# Sequential formation of Drosophila circuit asymmetry via prolonged structural plasticity

**DOI:** 10.1101/2025.05.27.656373

**Authors:** Johann Markovitsch, Daniel Mitić, Alisa Jiménez-García, Zane Alsberga, Sarah Kainz, Rashmit Kaur, Thomas Hummel

## Abstract

Structural and functional differences between brain hemispheres are a common feature of animal nervous systems with reduced bilateral asymmetry often linked to impaired cognitive performance. How neuronal left-right asymmetry is initiated and integrated into a bilaterally symmetrical ground pattern is poorly understood. Here we show that directional asymmetry of a Drosophila central brain circuit originates from axonal interactions of two classes of bilateral pioneer neurons. Subsequent recruitment of neighboring neuron classes into the asymmetric neuropil precursor results in hemisphere-specific microcircuits. Circuit lateralization requires dynamic expression of the cell adhesion molecule Fasciclin 2 to maintain structural plasticity in axonal remodeling. Reduced circuit asymmetry following cell type-specific Fasciclin 2 manipulation affect adult brain function. These results reveal an unexpected degree of developmental plasticity of late-born Drosophila neurons in the formation of a new circuit node via the lateralized recruitment of symmetric circuit components.

## Introduction

In bilaterian animals, the central nervous system (CNS) develops in an overall mirror symmetrical fashion, which is maintained in the adult neuronal organization. While it is widely assumed that bilateral CNS symmetry originally evolved due to selection for directed locomotion, increasing evidence indicates that the interhemispheric separation has been translated into functional differences between left and right homotopic regions. This phenomenon, called lateralization seems to have originated multiple times in parallel independently of brain size, indicating an increased fitness of neuronal left-right asymmetry^1,2^. Recent experimental data suggest that lateralization improves cognitive function,^3–5^ and a negative correlation of performance with reduced asymmetry^6–8^. Similarly, the link between information processing in the human brain and cerebral lateralization is supported by a reduction of left-right asymmetry in schizophrenia, dyslexia and autism spectrum disorders^9^.

The *Drosophila melanogaster* brain contains a bilateral synaptic neuropil, the Asymmetrical Bodies (AB)^6,10^, with left-right differences in volume^10^, neuronal composition^10^ and protein expression^6,8^. A reduction of AB asymmetry correlates with deficits in olfactory^6,8^ and courtship^8^ long-term memory, indicating a direct relevance of CNS lateralization for cognitive function. The AB is embedded in a series of midline neuropiles called the central complex (CX), that in addition encompasses the ellipsoid body (EB), the fan-shaped body (FB), the noduli (NO), the protocerebral bridge (PB)^10,11^. The CX defines an evolutionarily conserved structure of the insect and the crustacean brain^12^ with similarity in associated behaviors and conserved genetic developmental programs. Furthermore, the insect CX shares conserved structural and functional features with the vertebrate basal ganglia^13^. The AB circuit follows the grid-like organization of the CX^12,14,15^, in which small-field neurons project along the anterior-posterior axis and connect two or more neuropiles of the CX to form columns. These columns are crossed by perpendicular projections of large-field tangential neurons that connect the CX to extrinsic brain regions and provide the majority of synaptic input.

Although born as early as in the embryo^15^, CX neurons form synaptic connections not before pupa formation, making the CX functionally a structure of the adult organism^16^. Immature and bilaterally symmetric neuropile primordia composed of a tight bundle of undifferentiated neural processes and filopodia are part of the interhemispheric commissure of the early larval brain^15,16^, while lateralization of the CX occurs much later during pupal development^8,17^.The assembly of lateralized circuits requires the integration of left-right asymmetric circuits into the bilateral symmetric ground pattern and the adjustment of interhemispheric communication, a process poorly described so far.

Here, we show that the developmental interaction between two classes of pioneer afferent neurons initiate lateralized remodeling of the AB neuropile in which the dynamic expression of Fascilin2 controls the level of class-specific asymmetry. Lateralized AB circuit assembly within the symmetrical CX is mediated by the recruitment of early-born FB neurons to induce side-specific synapse formation. This level of structural plasticity in building neuronal connections across brain neuropils is rather unique at this advanced state of Drosophila adult nervous system development, revealing brain lateralization as an additional genetic program to reorganize the bilateral ground pattern.

## Results

### Two types of afferent neurons determine left-right asymmetry of Central Complex connectivity

The Asymmetrical Body (AB) is a bilateral neuropil at the ventral border of the fan-shaped body (FB, Fig. 1a,b). Its eponymous left-right asymmetrical neuronal organization is most evident in a different neuropil size between hemispheres with the right AB (AB-R) on average 4x larger in volume compared to AB-L^10^. In addition, the AB was originally identified in *Drosophila* due to its unilateral expression of the cell adhesion molecule Fasciclin 2 (Fas2) expression in AB-R^6^.

**Fig 1.**
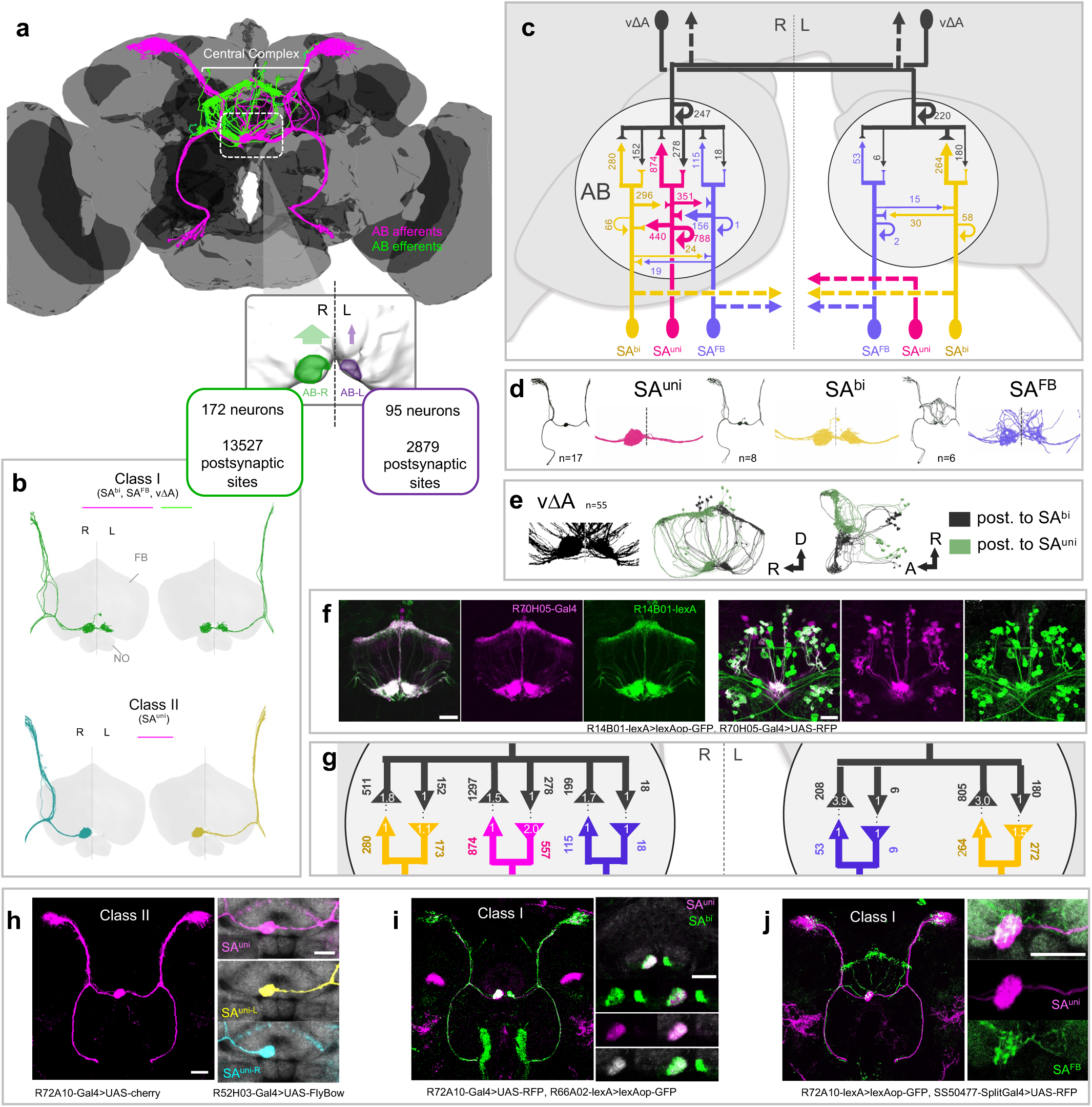
Interhemispheric connectivity of main AB neuronal types. **(a)** Position of the AB circuit in the adult Drosophila central brain (3D reconstruction from EM data, Flywire) and left-right asymmetry in synaptic connections in the AB neuropiles (EM data, Hemibrain). Main afferents in magenta, main efferents in green. **(b)** Two classes of morphological asymmetry of AB neurons: neurons display a bilateral connectivity pattern but differ in the innervation between AB-R and AB-L (Class I); neurons show unilateral connectivity to the same hemisphere (AB-R) (Class II). 3D reconstruction from EM data (Hemibrain). Fan-shaped body and noduli neuropiles in grey. **(c)** Schematic of the AB circuit and the number of presynaptic sites targeting the indicated type (arrowhead) in AB-R and AB-L (EM data, Hemibrain). **(d)** The 3D reconstruction from EM data (hemibrain) for 3 main AB afferent neurons types, the unilateral axon projection of SLP-AB-unilateral (SA^uni^) compared to the two bilateral AB classes SLP-AB-bilateral (SA^bi^) and SLP-AB-FB (SA^FB^). **(e,f)** 3D reconstruction of EM data (hemibrain) and expression lines for AB efferent neurons vΔA, which connect the AB with the dorsal FB. Immunofluorescence detection of lexAop-GFP driven by R14B01-lexA (used to label vΔA early pupal development, green) and UAS-RFP driven by R70H05-Gal4 (magenta). **(g)** Numbers and ratios of presynaptic sites and connected postsynaptic sites for SA^uni^ (magenta), SA^bi^ (yellow), SA^FB^ (blue) and vΔA (dark grey) in AB-R and AB-L. **(h)** Afferent SA^uni^ projections are hemisphere-dependent (Class II asymmetry): ipsilateral SA^uni-R^ and contralateral SA^uni-L^. Immunofluorescence detection of UAS-cherry, UAS-Flybow1.1 driven by R72A10- and R52H03-Gal4 and CadN antibody staining (grey). **(i)** Afferent SA^bi^ neurons (green) project bilaterally to AB-R and AB-L (Class I asymmetry). Fas2 expression (grey) fully overlaps with SA^uni^ in AB-R, but is absent in SAbi innervations of the dorsal AB-R and in AB-L. Immunofluorescence detection of UAS-RFP driven by R72A10-Gal4 (SAuni, magenta), lexAop-GFP driven by R66A02-lexA (SA^bi^, green) and anti-Fas2 antibody staining (grey). **(j)** Bilateral afferent SA^FB^ neurons (green) project both to the dorsal FB and AB-R/AB-L. Immunofluorescence detection of lexAop-GFP driven by R72A10-lexA (SA^uni^, magenta), UAS-RFP driven by SS50477-Split-Gal4 (SA^FB^, green) and CadN antibody staining. All scale bars, 20µm.

The AB neuropil is highly interconnected with the FB and the “Superior Lateral Protocerebrum” (SLP) in the dorsal adult *Drosophila* brain^18^ (Fig. 1a). Three classes of AB afferent neurons, SLP-AB-unilateral (SA^uni^/SA1&2), SLP-AB-bilateral (SA^bi^/SA3) and SLP-AB-FB (SA^FB^/SAF), provide tangential synaptic input into the AB neuropil (Fig.1d,h-j, also see methods for nomenclature), while columnar output neurons (vΔA) connect the AB to the dorsal FB^18^ (Fig.1e,f).

The cell bodies of all SA neuron classes colocalize in an anterior-ventral cluster and project in common tract to the dorsal SLP region (Fig. 1a,d). The SA projections extend a horizontal axonal branch to the ventral border of the CX to arborize in the AB neuropil (Fig. 1A, D-F). Compartment-specific marker expression (Syt::GFP, Brp:GFP DenMark::RFP) indicates a polarized SA neuron organization with the main dendritic input region in the SLP and the AB neuropil as the presynaptic axonal domain (Fig. 1). While their dendritic domains mostly overlap in the SLP region, SA^uni^, SA^bi^ and SA^FB^ afferent neurons differ in their presynaptic innervation between the AB neuropil in the right versus left hemisphere.

The axonal processes of SA^uni^ are the central node of connectivity in AB-R (Fig. 1c,g). Due to the larger number of cells compared to other AB afferents (about nine SA^uni^, four SA^bi^, three SA^FB^ per hemisphere), SA^uni^ define the dominant input neurons to vΔA, the main AB efferents. In addition, SA^uni^ is a central component of afferent-afferent interconnectivity, not only towards the other AB afferent neurons, in particular with SA^bi^ but also among neurons of the bilateral SA^uni^ groups (electron microscopy (EM) data^19^, Fig. 1c). Its capacity as a hub connecting efferent and afferent neurons is also reflected in the fact that not only presynaptic sites (Fig. S1a-a’’) but also about 60% of SA^uni^ postsynaptic sites are located in the AB neuropile (Fig. S1b-b’’), despite prominent dendritic branches in the dorsal SLP region. This interconnectivity between AB afferents and efferents appears to be specific to the SA^uni^ cell type, as the afferent SA^FB^ and SA^bi^ neurons form only sparse synapses with each other, both in AB-R but also in AB-L, where SA^uni^ processes are absent (Fig. 1c). This suggests a distinct ability of SA^uni^ to recruit synapse formation with other AB afferent neurons.

SA^uni^ bilateral asymmetry is unique among all neurons in the *D. melanogaster* brain, as this cell type in both hemispheres provides synaptic input only to AB-R via two morphological subtypes, the ipsilateral SA^uni-R^ and the contralateral SA^uni-L^ neurons (Fig.1b,h). While differences in the relative volume of afferent axon arborizations between AB-R and AB-L are found for all AB afferents, they do not develop hemisphere-specific projection patterns depending on cell body position (Fig.1b). To test if SA^uni-R^ and SA^uni-L^ neurons are intrinsically different in organizing AB lateralization we prevented contralateral projections across the midline in a hypomorphic *Neuroglian* mutant background (Nrg^849^), while leaving the overall SA^uni^ projections unaffected. Both SA^uni-R^ and SA^uni-L^ neurons innervate the ispsilateral AB neuropile and no directional size differences could be detected (Fig. 2). Induction of apoptosis in SA^uni^ had similar effects on vΔA volume (confirming results by Sakamura et al.^17^, Fig. 2e,e’). Furthermore, ipsilateral SA^uni-L^ neurons are able to form functional synapses with postsynaptic vΔA neurons (Fig. 2b’’’). These results indicate a critical role of SA^uni^ for the formation of overall CX left-right asymmetry. Development within separated hemispheres revealed no intrinsic differences between SA^uni-R^ and SA^uni-L^ in side-specific AB innervation and show how important interhemispheric communication is for lateralization.

**Fig 2.**
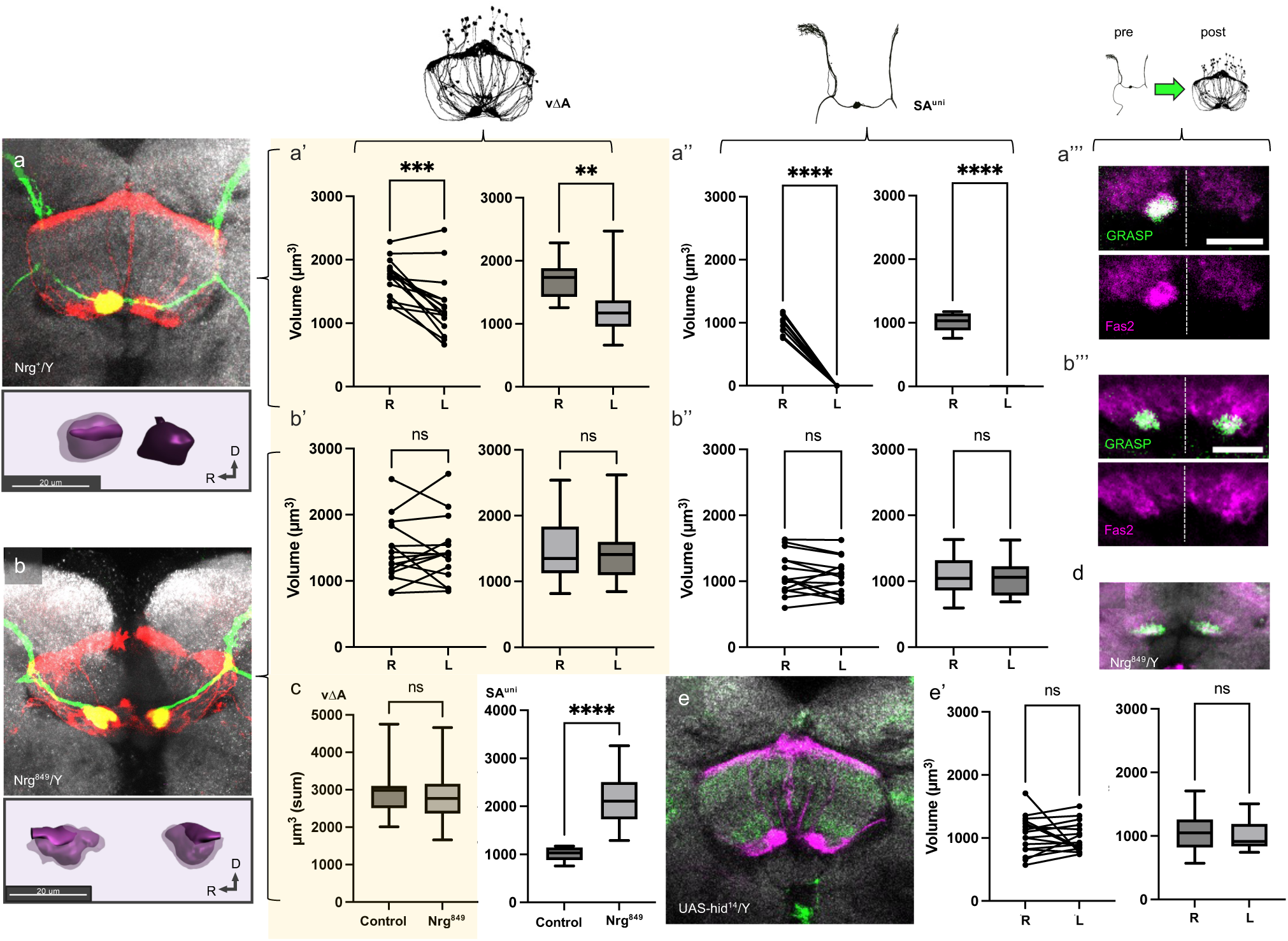
Developmental plasticity of lateralized AB innervation. **(a, b, e)** Bilateral SA^uni^ projection abolishes AB asymmetry. **(a, b, c, e)** Paired data plot (Wilcoxon Test for paired samples) to compare relationship of AB-R and AB-L measured from the same brain and boxplot to compare distribution of measured innervation volumes between AB-R and AB-L (Mann-Whitney U). In wild-type brains, the volume of innervations by vΔA neurons **(a’)** and SA^uni^ **(a’’)** is statistically significantly larger in AB-R than in AB-L (Mann-Whitney U). AB innervation volumes were measured from immunodetection of lexAop-GFP driven by R72A10-lexA (c) and UAS-RFP driven by R70H05-Gal4. **(a’’)** SA^uni^ neurons in all measured brains innervated only AB-R. **(a’’’)** Immunodetection of reconstituted GFP (green) from nysb-GRASP driven by R72A10-Gal4 (presynaptic) and R38D01-lexA (postsynaptic) and Fas2- (magenta) and CadN-antibody (grey) indicates functional synapses between SA^uni^ and vΔA in AB-R. **(b)** Hemizygosity for the hypomorphic allele Nrg^849^ prevents SA^uni^ neurons (as well as SA^bi^ **(d)**) from forming commissures across the midline, while decussating projections by vΔA neurons are not affected. The population-level volume difference of SA^uni^ and vΔA innervations between AB-R and AB-L that we found in the wild type was not detectable in the Nrg^849^ mutants analysed **(b’,b’’). (b’’)** GRASP showed functional and active synapses between Nrg-mutation-induced ipsilateral SA^uni^ neurons and vΔA in AB-R and AB-L. Immunodetection of reconstituted GFP (green) from nysb-GRASP driven by R72A10-Gal4 (presynaptic) and R38D01-lexA (postsynaptic) and Fas2- (magenta) and CadN-antibody (grey) staining. **(c)** The total AB volume innervated by SA^uni^ is twice as large in Nrg-induced ispislateral neurons compared to the control. **(d)** Immunofluorescence detection from lexA-GFP driven by R66A02-lexA and Fas2 (magenta) and CadN (grey) antibody staining. **(e)** vΔA innervation pattern in the central complex after ablation of SA^uni^ by expression of UAS-hid^14^ driven by R72A10-Gal4. Immunodetection of lexA-opGFP (magenta) driven by R70H05-lexA (vΔA) and CadN antibody staining (grey). **(e’)** Neither did the sign of the slope between the AB-R and AB-L volumes show a clear trend across individuals, nor did we find a statistically significant difference between the grouped AB-R and AB-L volumes. Significance threshold are: *<0.05; **<0.01; ***<0.001; ****<0.001.

With the absence of SA^uni^ axon terminals in AB-L the question arises as to how afferent neurons arrange their connections in the smaller neuropile. Compared to AB-R, AB-L generally contains fewer, but still a considerable number of synapses (Fig 1c,g). The presynaptic sites of the second major afferent class SA^bi^ on vΔA, appear to be evenly distributed between AB-L and AB-R (Fig. 1c). Interestingly, while both SA^FB^ and vΔA neurons develop less postsynaptic sites in AB-L compared to AB-R, vΔA redistributes its pool of postsynaptic sites to different synaptic partners, in particular SA^bi^ and other vΔA neurons (Fig. 1c and S1c-e’). As a result, synaptic weight of single SA^bi^ presynaptic sites is higher in AB-L than in AB-R (Fig. 1g), resulting in an opposite direction of asymmetry (AB-R<AB-L) with roughly double the number of vΔA postsynaptic sites in the left hemisphere. Thus, the right dorsal FB in AB receives predominantly input from SA^uni^ neurons, while the left dorsal FB receives input from SA^bi^ (Fig. 1f). While SA^bi^ in AB-L interacts directly with vΔA, in AB-R it equally modulates the SA^uni^-ΔA input to vΔA, thus providing direct and indirect input to vΔA. This very likely translates the structural asymmetry of the AB circuit into asymmetry at the activity level in the CX.

In summary, while SA^uni-R^ and SA^uni-L^ seem intrinsically identical to innervate the ipsilateral AB neuropile, the presence or absence of this AB afferent class due to its unique ipsi-/contra-lateral axon morphology translates into distinct microcircuit organization: a prominent recurrent afferent-afferent connectivity in AB-R and a feed-forward afferent-efferent organization in AB-L common to other CX circuit motifs^20^.

### Cellular interactions during lateralized circuit remodeling

This lack of input asymmetry between AB-L and AB-R distinguishes SA^bi^ connectivity from SA^uni^, although the two neuron types appear morphologically almost identical, especially in the sporadic bilateral phenotype/variant of SA^uni^. In this case, which develops in 7.6% of wild-type flies^6^, SA^uni-L^ and SA^uni-R^ both project to AB-L and AB-R, but maintain a bilateral asymmetry in axon arborization towards AB-R (Fig. 5a). We found SA^bi^ neurons (labelled by R66A02-Gal4/-lexA) present in all brains bilaterally innervating AB-L and AB-R independent of SA^uni^ projection pattern (Fig.1i). While both uni- and bilateral SA^uni^ morphotypes express the cell-adhesion molecule Fas2, adult SA^bi^ neuronal processes in AB-R did not fully overlap with Fas2 antibody staining and were clearly Fas2-negative in AB-L (Fig.1i). These results confirm the hemibrain annotation of SA^uni^ and SA^bi^ as two AB afferent neuron types with unique morphological and molecular features.

The distinct expression of Fas2 in SA^uni^ and the reported role of Fas2 in synaptic maturation and remodeling suggest that Fas2 may be involved in SA^uni^-induced lateralized connectivity. To test this, we used RNAi to suppress Fas2 expression under the control of endogenous Fas2 enhancer regions (Fas2^Mz507^-Gal4, Fas2^MiMIC12989^-Gal4), which resulted in Fas2 no longer being detectable in the CX with antibody staining. Developmental loss of Fas2 results in a significant increase in bilateral SA^uni^ neurons up to 40% (Fig. 5a), indicating a role of Fas2 expression in CX lateralization not only at the synaptic level but also at the level of cell morphology.

We next analyzed Fas2 expression during CX neuropile formation via monoclonal antibody labeling of all transmembrane isoforms (1D4^21^) as well as an endogenously GFP-tagged Fas2 allele (Fas2^GFP397^). Throughout pupal development we observed a close overlap between Fas2 antibody labeling and Fas2^GFP^ fluorescence signals, indicating the expression of mostly transmembrane isoforms in the developing central complex (Fig. 3a). At 12h after puparium formation (APF), a thin bundle of Fas2- positive commissural fibers at the ventral surface of the CX primordia could be detected, which increased in thickness until 20h APF, when a bilateral pair of AB precursors started to segregate as distinct neuropils (Fig. S2). At 25h APF, no consistent directional lateralization of Fas2 expression was evident but small fluctuating asymmetry in neuropil size could be observed among wild type brains (Fig. 3c). Over the next 10 hours a relative volume increase of aggregated Fas2-positive neuronal processes in the right hemisphere accompanied by an increase in Fas2-reporter expression could be observed. At 36h APF the right AB precursor was typically not only larger in size but also had stronger Fas2 expression levels. In addition, an enriched Fas2 expression in the ventral domain of the AB-R neuropile compared to the dorsal domain could be detected. In the majority of analyzed brains Fas2 signal in AB- L disappears around 40h APF with rare extended expression until 45h APF (Fig. 3a).

**Fig 3.**
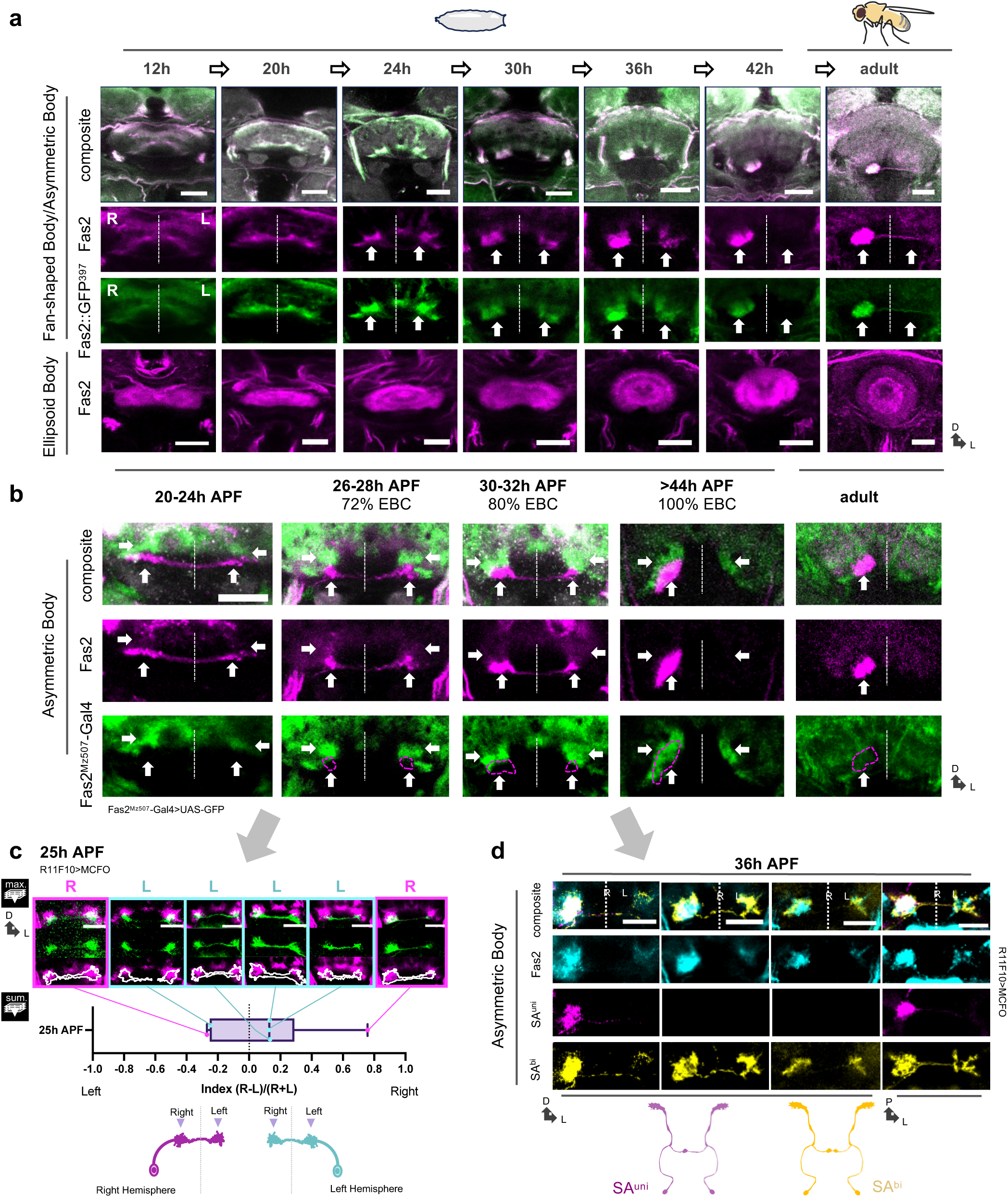
Fas2 is dynamically expressed in AB development. **(a)** Pupa brains dissected at the indicated number of hours after puparium formation (h APF). Immunodetection of Fas2::GFP^397^ (green) and Fas2 (magenta) and CadN (grey) antibody staining. **(b)** Fas2^Mz507^-Gal4, that drives Gal4 expression under control of Fas2 regulatory regions, did not drive expression in the ventral AB primordium and was not expressed in the adult AB, although the ventral and dorsal primordium and the adult right neuropile were labelled by Fas2 antibody staining. Immunofluorescence detection of UAS-GFP driven by Fas2^Mz507^-Gal4 (green) and Fas2 (magenta) and CadN (grey) antibody staining. Arrows indicate position of dorsal and ventral AB-R and AB-L primordia. **(c)** Individual cell clones innervating the ventral AB primordium at 25h APF were not yet uniformly lateralised. Regardless of the position of the cell body (R: right hemisphere, L: left hemisphere), neurons tended to be more densely innervated in AB-R compared to AB-L. Immunofluorescence detection from MCFO (HA or FLAG) driven by R11F10-Gal4 (green) and Fas2 antibody staining (magenta). **(d)** SA^uni^ occupies the ventral, SA^bi^ the dorsal AB primordium. Immunofluorescence detection from MCFO (HA or FLAG) driven by R11F10-Gal4 (magenta: clones segregated to the ventral primordium, yellow: clones in the dorsal primordium) and Fas2 antibody staining (cyan). All scale bars, 20µm.

Next, we compared the expression of the two Fas2 Gal4 driver lines which induces a bilateral SA^uni^ phenotype, Fas2^Mz507-^Gal4 (Fig. 3b & S3a) and Fas2^MiMIC12989^-Gal4 (Fig. S3b) with Fas2 protein expression. Interestingly, neither Fas2 Gal4 lines drove reporter expression in the adult AB. Upon expression of UAS-mCD8::GFP during pupal development, both lines labeled neurons that innervate only in the dorsal AB precursors, while both dorsal and ventral area are Fas2-positive, as demonstrated by antibody staining. Decline of Gal4 expression coincided with Fas2 protein no longer detectable in AB-L. These results indicate that the early AB neuropile is formed by two moleculary-distinct populations of afferent neurons, which segregate their axons along the dorso-ventral axis of the AB precursors.

To match the Fas2-positive neurons in AB development with adult morphological afferent types, we analysed MCFO mosaics using R72A10- (SA^uni^), R66A02- (SA^bi)^ and R11F10-Gal4 (SA^uni^ and SA^bi^) lines (Fig. 3a,d & S6). While both SA^uni^ and SA^bi^ cell types formed bilateral axonal projections in early pupa stages, SA^uni^ neurons clustered to ventral AB-L and AB-R subregions, while SA^bi^ densely occupied the dorsal AB neuropils (Fig. 3d, S6). These dorsal and ventral axon domains mostly merge in subsequent AB maturation, but SA^bi^ processes could still be found more densely packed dorsally in the adult AB-R. With the described birth-order dependent dorsal-ventral innervation of FB layers by LAL-derived tangential neurons^22^, SA^bi^ neurons would be generated shortly before SA^uni^ as the last cell identity to emerge from the LALv1A hemi-lineage. In addition to the late-born SA^uni^ and SA^bi^ classes, a third class of AB afferent neurons, SA^FB^ (SS50477-SplitGal4) emerges within the LALv1A hemi-lineage several divisions^22^ before SA^uni^ and SA^bi^ and project towards the central complex. In contrast to SA^uni^ and SA^bi^, SA^FB^ axons bypass the future AB precursor region and extended to the dorsal FB (Fig. 4a). Thus, the initial formation and the asymmetric remodeling of the AB neuropiles occurs through axonal interactions between the sequentially-born^22^ SA^bi^ and SA^uni^ pioneer afferents.

**Fig 4.**
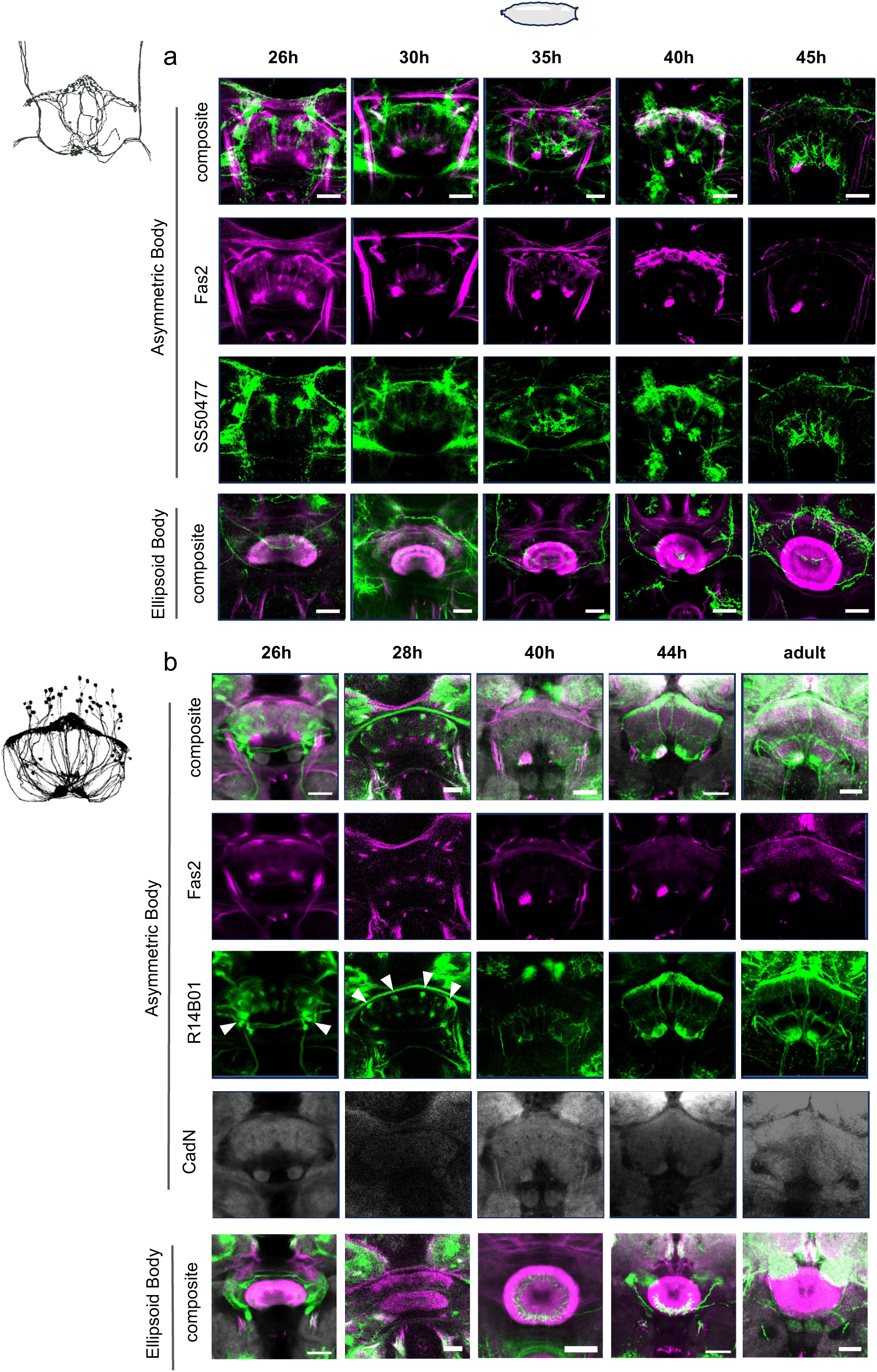
SA^FB^ afferents and vΔA output neurons project to the FB early, but innervate AB-R and AB-L only after SA^uni^ and SA^bi^ remodelling is complete. **(a)** Immunodetection from UAS-Flag driven by SS50477-Split-Gal4 (green) and Fas2 antibody staining (maganta). **(b)** vΔA projections arrive in dorsal and ventral FB before 30h APF (arrowheads). Immunodetection from UAS-GFP driven by R14B01-Gal4 (green) and Fas2 antibody staining (maganta). All scale bars, 20µm.

At 25h APF, individual SA^uni^/SA^bi^ afferent neurons started to increase axonal branching in AB-R accompanied by denser distal innervations. However, initial axon branch extension was rather dynamic and no clear SA^uni^ lateralization could be recognized at this time point (Fig. S6b). From 25h APF to the completion of lateralization at 36h APF, the SA^uni^ axonal processes underwent a considerable remodeling, withdrawing from AB-L and restricting their innervation exclusively to AB-R. Axonal processes of single SA^uni^ neurons were not labeled by Fas2 antibody staining, suggesting that the onset of expression started during axonal remodeling or could be explained by strict localization of Fas2 protein within the AB neuropil. With the retraction of SA^uni^ axons from AB-L, the closely associated SA^bi^ axons also developed asymmetrically with denser innervations in AB-R, but sparse axonal processes remained at AB-L after the completion of SA^uni^ remodeling (Fig. 3d). SA^bi^ neurons expressed Fas2 throughout development, but gradually attenuated expression after 36h APF (Fig. 3d & S6a). The loss of Fas2 expression in SA^bi^ at 40h APF indicated the completion of SA^uni^ and SA^bi^ axon remodeling. Interestingly, at the same time ventral processes of SA^FB^ neurons start to innervate the asymmetric AB neuropile in both hemispheres (Fig. S6 & 4a).

As with SA^FB^ afferents, no innervation of the bilateral AB neuropil by processes of vΔA neurons, the main AB efferent class, could be observed before 40h APF. vΔA neurons, initially targeted the dorsal and ventral FB during early pupal development and form restricted Fas2-negative fiber aggregates by 30h APF, reminiscent of early-born pontine neurons of the FB^15,23^. No innervation of the bilateral AB neuropil by vΔA processes could be observed before the completion of the unilateral SA^uni^ axon remodeling. Extending dendritic processes of vΔA neurons show a different growth pattern between the left/right AB primordia, covering loosely the outer surface of AB-R while tightly converging in the center of AB-L (Fig. 4b). Of note, about 30% of vΔA neurons never invade the AB. Instead they retain the initial pontine projection pattern into the adult system connecting FB layer 1 instead of AB-R or AB-L to layers 8 and 9 (hemibrain data^18^). vΔA cell bodies form four spatially separated clusters per hemisphere, which reflect their descent from the cell lineages DM1-4^24–26^. Their fibers follow the described projection pattern^15^, with the vΔA neurons derived from DM1 and DM2 projecting contralaterally and the vΔA neurons derived from DM3 and DM4 projecting mainly ipsilaterally with respect to the position of their cell bodies (Fig. S5). The majority of individual vΔA neurons innervate only AB-R or AB-L in both the hemibrain dataset and our mosaic analysis, with a small fraction (of all 4 lineages) display a bilateral AB-R/L connectivity via the commissural tract of SA afferents (Fig. S1e). Interestingly, EM data revealed a left-right asymmetry in synaptic differentiation of these bilaterally innervating vΔA neurons with increase postsynaptic sites in AB-R (Fig. S1e’), indicating a higher level of morphological plasticity compared to the stereotype pattern of CX principal neurons.

To further determine the morphological variability of the AB circuit we analyzed the projections of LCNOpm, which are among the first born neurons of the LAL lineage^22^. LCNOpm neurons connect the Noduli and LAL neuropiles with AB-R but not AB-L^18^, thereby defining a second class of bilaterally asymmetric neurons in the *D. melanogaster* CX (Fig. S4). Class-specific expression lines (SS47384, SS47392) revealed a high level of variability in the amount and pattern of AB innervation by LCNOpm neurons in the right hemisphere: Significant levels of AB innervation was only detected in 17 % of brains analysed, while in the majority of brains (44.5 %) LCNOpm neurons extend in the direction of the ABs but terminate before, without arborizations (Fig. S4b-b’’’). Interestingly, similar to SA^uni^ neurons, this lateralization towards the right AB neuropil is strongly reduced in Nrg mutants with multiple cases of LCNOpm neurons in the left hemisphere also show additional AB innervation.

In summary, the data presented here indicates a robust developmental program for AB neuropile formation and axon guidance which involves afferent-afferent interactions while the synaptic recruitment of output neurons occurs after neuropil lateralization is complete. Secondary integrated AB neuron classes exhibit different degrees of stochasticity in the adult level of lateralized neuropil innervation.

### Dynamic Fas2 expression supports afferent lateralization independent of cell adhesion

With dynamic Fas2 expression in developing AB afferents and reduced lateralization phenotypes following Fas2 reduction, we next used targeted RNAi knockdown with cell-type specific drivers to identify the neuronal identity of critical Fas2 expression (Fig. 5b). Since the AB-R specific innervation pattern of SA^uni^ neurons correlates with CX lateralization in the adult CX (see chapter 1), we visualized their L/R innervation pattern (R7210-lexA/LexOpGFP) following the targeted manipulation of Fas2 expression. Loss of Fas2 in SA^uni^ (R72A10) reduced Fas2 expression in the adult AB-R, but the unilateral restriction of SA^uni^ processes appeared unaffected (confirming results by Sakamura et al.^17^). In contrast, knockdown of Fas2 in SA^bi^ (R66A02-Gal4) was able to replicate the effect of the manipulation under the control of Fas2^Mz507^-Gal4. As indicated by single cell clones, SA^bi^ neurons aggregate during development in the dorsal AB subregion, which correlates with the Fas2^Mz507^-Gal4 expression, but only transiently express Fas2 protein.

**Fig 5.**
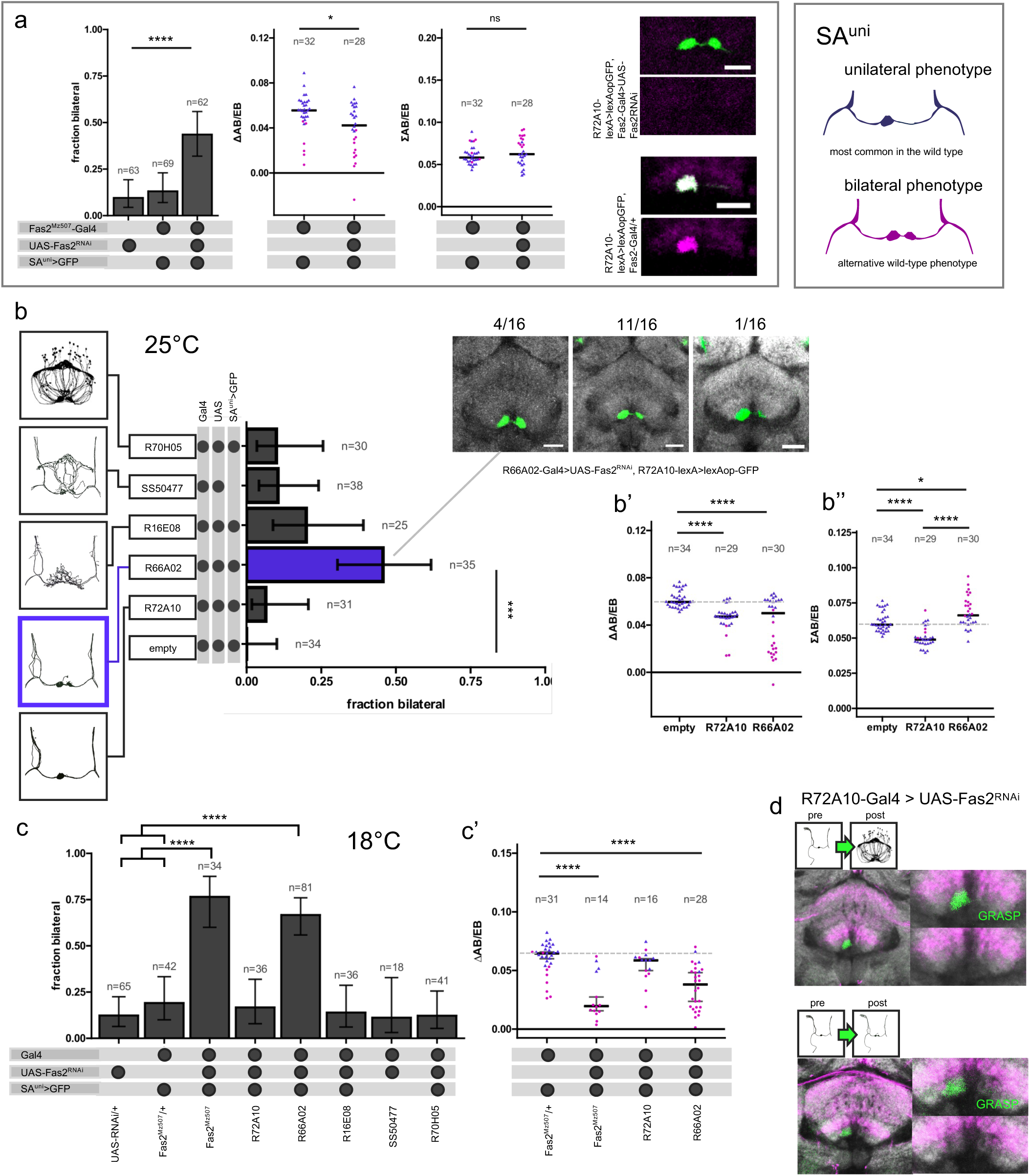
Fas2 function supports central complex lateralization. **(a)** Targeted RNAi knockdown of Fas2 driven by Fas2^Mz507^-Gal4 at 25°C increased the rate of brains with SA^uni^ projecting bilaterally to AB-R and AB-L (χ^2^) decreased innervation area lateralization (ΔAB/EB, Mann-Whitney U). Although Fas2^Mz507^-Gal4 does not drive expression in SA^uni^ directly, Fas2 antibody staining was no longer detectable in the AB. Immunofluorescence detection from lexAop-GFP (green) driven by R72A10-lexA and Fas2 antibody staining (magenta). **(b)** Knockdown driven by driver lines cell-type specific for AB neurons showed that only the loss of function in SA^bi^-specific R66A02-Gal4 has a similar effect as loss in Fas2^Mz507^-Gal4-positive cells on SA^uni^ projection pattern (χ^2^). **(b’)** Loss of Fas2 reduced SA^uni^ lateralization Kruskal-Wallis/Dunn) cell-autonomously (R72A10-GAL4) and non-cell-autonomously (R66A02-GAL4). **(b’’)** The bilateral brains had higher total innervation area, resulting in a higher area for R66A02-Gal4 experimental group, while loss of Fas2 in SA^uni^ reduced innervation area without affecting the projection pattern (Kruskal-Wallis/Dunn). **(c)** Targeted RNAi knockdown of Fas2 at 18°C developmental temperature increased the the expressivity of the bilateral SA^uni^ phenotype (χ^2^) and **(c’)** reduced lateralization when driven by Fas2^Mz507^-Gal4 or R66A02-Gal4 (Kruskal-Wallis/Dunn). **(d)** Immunofluorescence detection from nsybGRASP (green) indicates the formation and maintenance of functional synapses by SA^uni^ in the absence of cell-autonomous Fas2 expression. Fas2 (magenta) and CadN (grey) antibody staining. Blue triangles indicate measurements from brains with unilateral AB projection by SA^uni^, magenta dots indicate brains with bilateral projections. Crossbars indicate medians. Error bars indicate two-sided 95% confidence intervals for the single proportions. Significance threshold are: *<0.05; **<0.01; ***<0.001; ****<0.001.

To further investigate a potential role of Fas2-mediated cell adhesion in CX lateralization, we expressed different Fas2 isoforms (UAS-Fas2^PEST+^ and UAS-Fas2^PEST-^) in afferent SA^uni^ and SA^bi^ neurons (Fig. 6). Overexpression of transmembrane isoforms of Fas2 (UAS-Fas2^PEST+^ or UAS-Fas2^PEST-^) in SA^bi^ axons (Fas2^Mz507^-Gal4 and R66A02-Gal4) strongly impairs asymmetric remodeling of SA^uni^ processes (∼75% of brains showing bilateral innervation of SA^uni^ neurons, Fig. 6a). In contrast, overexpression of just the intracellular or extracellular domains of Fas2 (UAS-intra-Fas2^PEST-^::YFP and UAS-extra-Fas2^PEST-^ ::YFP^27^) under control of Fas2^Mz507^-Gal4 did not affect lateralization (Fig.6b).

**Fig 6.**
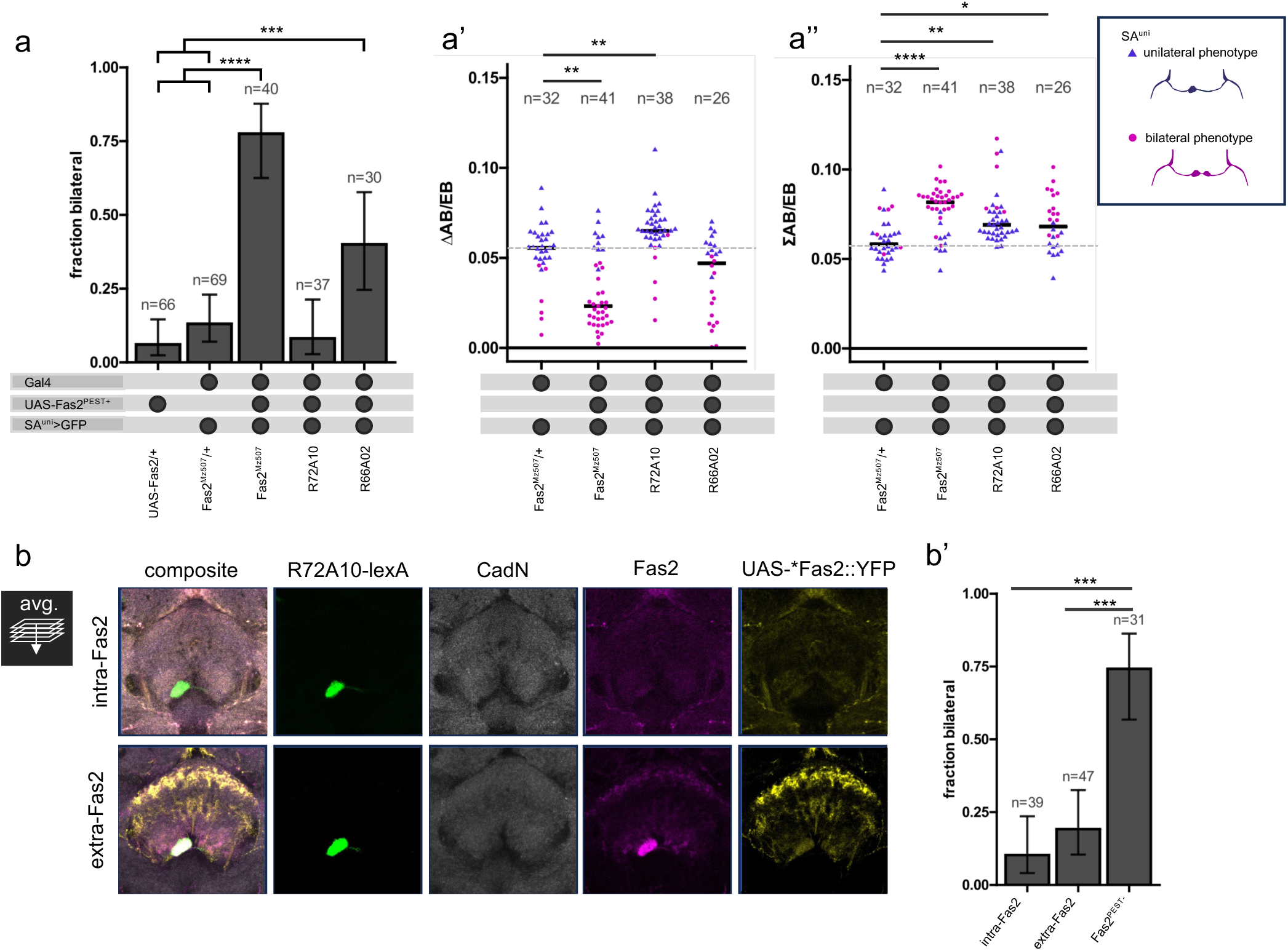
Overexpression of Fas2 transmembrane isoforms in SA^bi^ reduces CX lateralization while increasing AB innervation by SA^uni^. **(a)** Overexpression of the Fas2^PEST+^ isoform in drivers that label SA^bi^ (R66A02-Gal4 and Fas2^Mz507^-Gal4) increases the rate of brains with bilateral SA^uni^ projections (χ^2^) and decreases innervation area lateralization (Kruskall-Wallis/Dunn) **(a’). (a’’)** The normalized AB innervation area of SA^uni^ is increased in all Fas2 overexpression experiments (Kruskall-Wallis/Dunn). Blue triangles indicate measurments from brains with unilateral AB projection by SA^uni^, magenta dots indicate brains with bilateral projections. **(b)** Expression of UAS-intra-Fas2^PEST-^::YFP under regulatory control of Fas2^Mz507^-Gal4 results in a downregulation of endogenous Fas2 (magenta) in the central complex (AB-R, ventral and dorsal FB, inner EB). Expression of UAS-extra-Fas2^PEST-^::YFP under regulatory control of Fas2^Mz507^-Gal4 results in ectopic expression of endogenous Fas2 in FB.l1 & -.l8. No downregulation of Fas2 expression was observed. Immunofluorescence detection of intra-/extra-Fas2::YFP driven by Fas2^Mz507^-Gal4 (yellow), lexAop-GFP driven by R72A10-lexA, Fas2 (D4) antibody staining (magenta) and CadN antibody staining (grey). **(b’)** Truncated Fas2 isoforms under control of Fas2^Mz507^-Gal4 do not recapitulate the expressivity of SA^uni^ projections of the full-length transmembrane Fas2 (χ^2^). Crossbars indicate medians. Error bars indicate two-sided 95% confidence intervals for the single proportions. Significance threshold are: *<0.05; **<0.01; ***<0.001; ****<0.001.

Similar to the knock-down experiment, the overexpression of Fas2 in SA^uni^ (R72A10-Gal4) had no effect on the differentiation of SA^uni-L^ and SA^uni-R^ (Fig. 6a). This data suggests a non-autonomous function of Fas2 in SA^uni^ remodeling (independent of SA^uni^-SA^bi^ trans-neuronal adhesion). Furthermore, the extracellular domain alone was not able to recapitulate the overexpression effect that we observed with the full transmembrane isoforms. To test whether Fas2 function is restricted to the interactions between the afferents SA^uni^ and SA^bi^, we expressed Fas2^RNAi^ in the FB neurons vΔA (R70H05-Gal4) and SA^FB^ (SS50477). Knockdown of Fas2 in vΔA and SA^FB^ had no effect on AB lateralization (Fig. 5a), consistent with our observation that these cell types innervate the AB after SA^uni^ and SA^bi^ remodeling is complete. These results reveal a critical role of SA^uni^-SA^bi^ cellular interaction in CX lateralization, in which the bilateral SA^bi^ support SA^uni^ axon remodeling.

### Fas2 delays afferent-afferent synapse formation

As the loss of Fas2 in developing SA^uni^ allow their axons still to aggregate at the positions comparable to the wild-type AB and to segregated from the adjacent CX neuropiles (CadN antibody staining), Fas2 function is not necessary for afferent axon guidance and bilateral AB neuropil formation (Fig. 5a). Using the nsyb-GRASP construct, we were able to confirm the presence of functional synapses in the adult AB circuit even in the absence of Fas2 (Fig. 5d), suggesting that connections were not eliminated as described for synapses in the NMJ^28^. While in Fas2-mediated cell adhesion has to be weakened to allow pruning of mushroom body γ neurons^29^, a similar role of Fas2 in the remodeling of SA^uni^ axons is not supported by our data, since neither overexpression nor knockdown of Fas2 affected their unilateral restriction.

We next examined a potential role of Fas2 in the synaptic growth within the AB circuit, as Fas2 has been described to modulate number of presynaptic sites dependent on expression level^28^. We measured the area of SA^uni^ axonal innervations in the ABs from our UAS-Fas2^PEST+^ and UAS-Fas2^RNAi^ data (Fig. 5a,b’,b’’ and Fig. 6a’,a’’). When under control of R72A10-Gal4, normalized area of AB innervations by SA^uni^ was decreased in the Fas2 knockdown condition and increased by Fas2 overexpression, thus affecting CX input lateralization without altering wild-type projection pattern. This supports Fas2 mediated synaptic growth in the AB circuit and indicates a cell-autonomous function of Fas2 in SA^uni^. While bilateral projections of SA^uni^ typically lead to smaller innervations in AB-R, the sum of the innervation cross section areas in AB-R and AB-L in the bilateral phenotype (both in wild-type variations and in experimental groups) tends to be larger than the area in the unilateral AB-R morphology (Fig. 5 and 6), indicating additional ectopic synapses of bilateral SA^uni^ projections. To test if the loss of Fas2 expression in SA^bi^ supports the formation of ectopic synaptic connections of SA^uni^, we prolonged AB remodeling via changes in environmental temperature. Previous studies have shown that reduced developmental temperature results in an increased formation of ectopic synapses during *Drosophila* pupal stages^30^. We compared the SA^uni^ asymmetry pattern as normalized difference between AB-R and AB-L following the targeted Fas2 knockdown in SA^bi^ at 18°C and 25°C. Here, the expressivity in the reduction of SA^uni^ asymmetry was higher at a developmental temperature of 18°C compared to 25°C (Fig. 5c,c’). Despite a reduced RNAi efficiency at lower temperatures, the effect increased significantly at 18°C compared to the experiment at 25°C (rank biserial, r=0.84 [95% CI: 0.69, 0.92], very large effect at 18°C; r=0.36 [95% CI: 0.08, 0.59], indicating a large effect at 25°C). These results support a model in which different level of Fas2 expression between closely-associated axon branches of SA^uni^ and SA^bi^ prevent inter-axonal synapse formation before the remodeling of AB pioneer afferents.

### SA^bi^ organizes CX lateralization

To further characterize the role of SA^bi^ neurons in supporting SA^uni^ lateralization we analyzed the previously described axon guidance molecule Netrin B (NetB), which signals repulsion via unc-5 receptor, but the cellular source of NetB is unknown^8^. Targeted knockdown of NetB in SA^bi^ driven by R66A02-Gal4 and Fas2^Mz507^-Gal4 resulted in a strong reduction of SA^uni^ remodeling compared to the control, while knockdown in developing SA^uni^ neurons (R72A10-Gal4) had no effect on AB lateralization (Fig. 7a). Interestingly, targeted knock-down of unc-5 in developing SA^bi^ also impact SA^uni^ remodeling (Fig. 7b), suggesting that the combined initial AB-L retraction of both afferent neurons (as described above) is critical for the complete SA^uni^ lateralization compared to the partial SA^uni^ retraction (type 2 versus type 1 asymmetry, respectively; Fig. 1b). This differential sensitivity to repulsive Netrin signaling is further supported by the loss of a remodeling phenotype for SA^uni^ following a temperature-mediated reduction in the level of unc-5 knock-down (Fig. 7c). SA^bi^ therefore needs to coordinate two critical developmental functions of CX lateralization: instructing the directed retraction of SA^uni^ axons via NetB secretion and preventing the formation of stabilizing synaptic connections before axon remodeling. Interestingly, although the total number of flies with bilateral SA^uni^ innervation (phenotypic penetrance) is higher in samples with SA^bi^-targeted NetB knockdown, the reduction of interhemispheric asymmetry (expressivity) in experimental brain with bilateral SA^uni^ innervation is stronger following the loss of Fas2 function in SA^bi^ (Fig. 5b & 7a) (ΔAB/EB, R66A02 vs. empty-Gal4, rank biserial: NetB^RNAi^: r=0.04 [95% CI: -0.23, 0.30], tiny effect vs. Fas2^RNAi^: r=0.29 [95% CI: 0.05, 0.49], indicating a medium effect). Of note, following the impairment of both, Fas2 or NetB function in SA^bi^, a reduction or loss of AB lateralization or even (but only in rare cases) a switch in the direction of AB asymmetry can be observed, indicating separate developmental processes.

**Fig 7.**
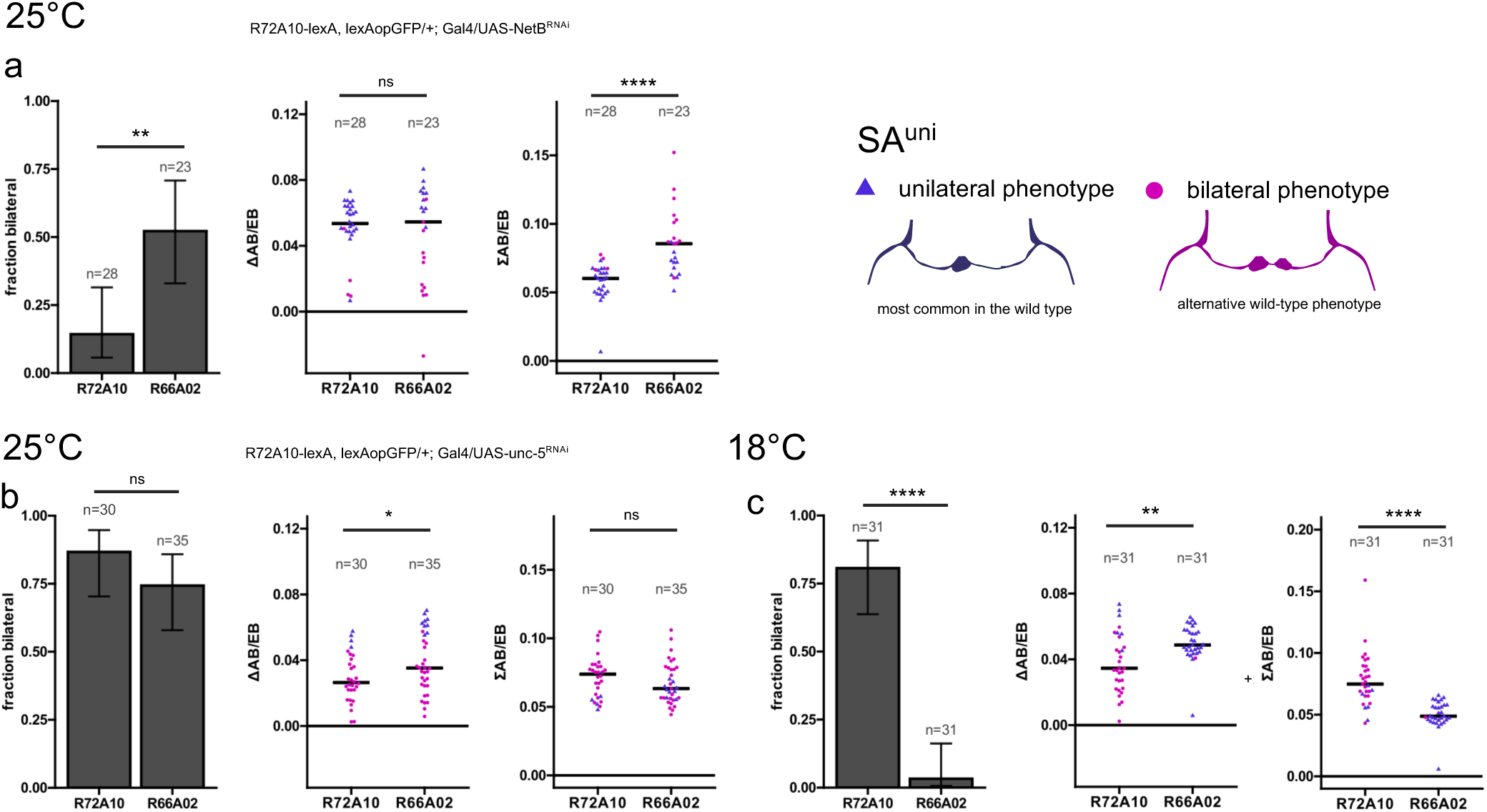
Netrin signalling proteins are expressed differently by SA^uni^ and SA^bi^. **(a)** At standard temperature, the rate of brains with SA^uni^ projecting bilaterally to AB-R and AB-L is statistically significantly higher when NetB^RNAi^ is expressed in SA^bi^ when compared to expression in SA^uni^ (χ^2^). While **t**he affect on input lateralization (ΔAB/EB) is low and not statistically significantly different between NetB^RNAi^ expressed in SA^uni^ or SA^bi^ **(a’)**, SA^uni^ has a larger AB innervation area (∑AB/EB) in a loss of NetB in SA^bi^ background (Mann-Whitney *U*) (**a’’**). **(b)** Loss of unc-5 receptor activity in SA^bi^ causes non-cell-autonously a comparable rate of bilaterally projecting SA^uni^ as when unc-5^RNAi^ is expressed in SA^uni^ itself (χ^2^), although the effect on SA^uni^ lateralization (ΔAB/EB) is weaker (Mann-Whitney *U*) **(b’)**. **(c)** In contrast to Fas2, developmental temperatures interacted differently with the expression in SA^uni^ or SA^bi^ . While there is no obvious difference in the cell-autonomous effect on SA^uni^ morphology (Mann-Whitney *U*), the non-cell-autonomous effect of unc-5 knockdown in SA^bi^ was no longer detectable (χ^2^). Innervation area was measured from SA^uni^ innervations in AB-R and AB-L. Error bars indicate two-sided 95% confidence intervals for the single proportions. Crossbars in area measurements indicate group medians. Blue triangles indicate measurements from brains with unilateral AB projection by SA^uni^, magenta dots indicate brains with bilateral projections. Significance threshold are: *<0.05; **<0.01; ***<0.001; ****<0.001.

As AB circuitry has been described to be involved in olfactory learning^31^ and satiety-dependent sugar preference^32^, with lateralization of SA^uni^ in particular being critical for aversive olfactory and courtship long-term memory^6,8^, we wanted to test if manipulation of bilateral SA^bi^ neurons is sufficient for memory impairment. We found that 24h memory expression in flies trained in a single-session appetitive olfactory conditioning paradigm^33^ was significantly reduced following Fas2 overexpressing in SA^bi^ (UAS-Fas2^PEST+^ under control of R66A02-Gal4, Fig. S7d). Similarly, post-fasting fructose preference was lost in flies with SA^bi^-induced reduction in AB lateralization (Fig. S7b-b’’) supporting the importance of developmental AB afferent interactions for efficient information processing in the mature brain.

## Discussion

Despite its omnipresence in nervous systems throughout the animal kingdom to support cognitive performance and behavioural plasticity, little is known of how brain lateralization is organized and established within neural circuits. As structural and functional differences of brain circuits between the right and left hemisphere are deeply embedded in an overall bilaterally symmetric nervous system, understanding the developmental programs that induce local symmetry breaks without interfering with symmetric pattern formation will provide important insights into the function and dysfunction of brain laterality. Using the *Drosophila* central complex as a conserved central brain structure in which a multitude of tangential and columnar neuron classes integrate in a highly symmetric bilateral circuit structure, we show how the interaction of two bilateral afferent neurons creates a local asymmetry followed by the synaptic integration and secondary lateralization of default symmetric CX neurons.

Lateralization of the CX depends on the interhemispheric interaction of two classes of AB pioneer neurons, SA^uni^ and SA^bi^, which not only initiate the formation of the AB primordia but also regulate the remodeling of a bilaterally symmetric ground state. Following unilateral retraction of SA^uni^ axons, earlier-born FB neurons are recruited into the AB primordium, including both SA^FB^ tangential neurons of the same lineage but also columnar vΔ neurons of the conserved DM lineages. SA^bi^ neurons instruct SA^uni^ lateralization via the cell surface molecule Fascilin 2 and the secreted guidance factor Netrin-B. The adult circuit connectivity depends on the SA^uni^ projection pattern. Although both AB afferent neurons derive from the same temporal window of the vLAL1A lineage, the SA^uni^ cell fate is specified in the last 8-10 final NB divisions, which might result in distinct differentiation states of the 2 afferent neurons by the onset of axon remodeling. Interestingly, while both cell types start to retract their axons from AB-L, SA^bi^ neurons maintain some processes in the bilateral ground state and thereby differ from the unilateral innervation of their sister neurons. From this developmental perspective, the bilateral morphotype of SA^uni^ can be explained by the incomplete segregation from their SA^bi^ sister neurons (see model in Fig. 8).

**Fig 8.**
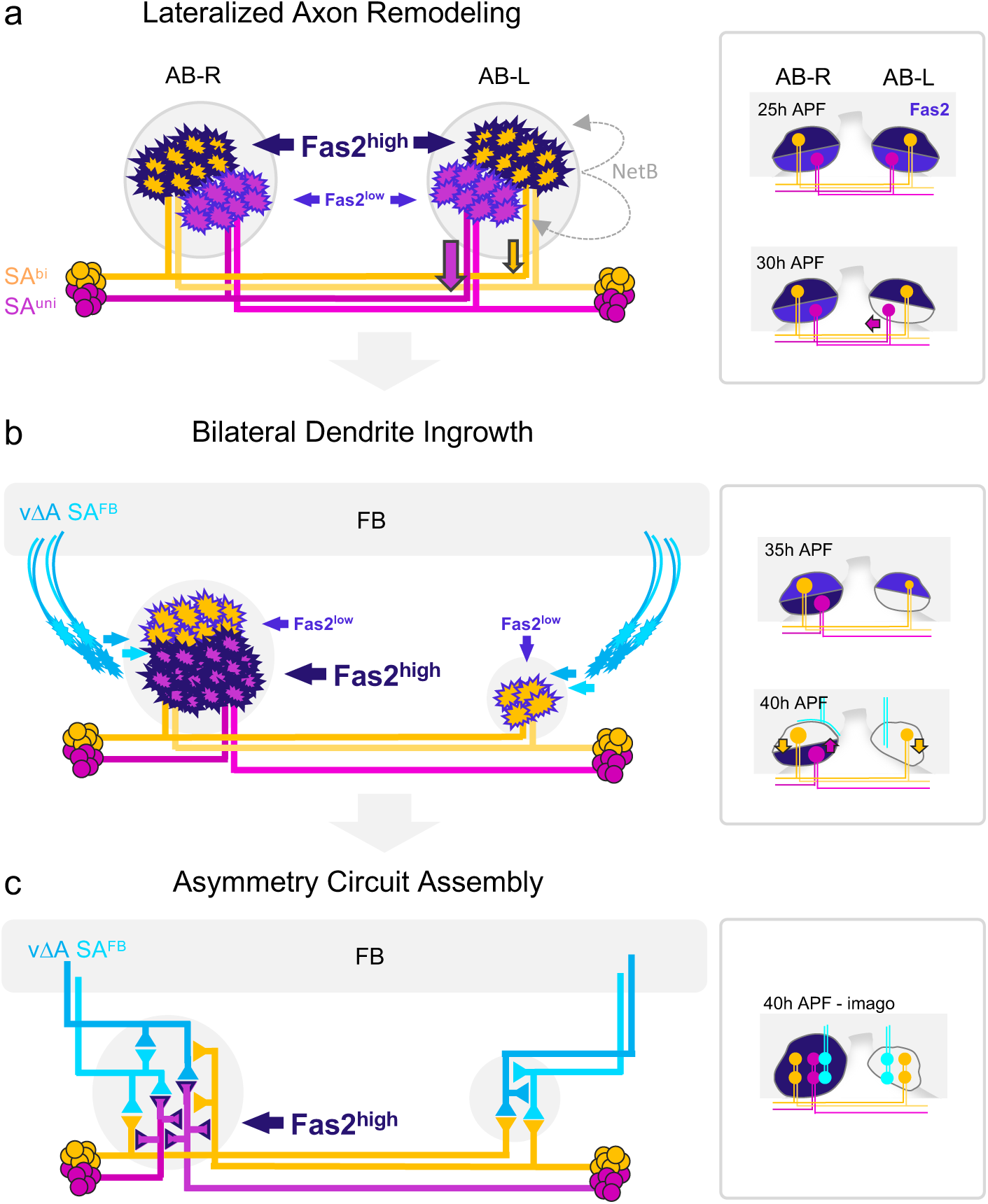
Sequential formation of Drosophila circuit asymmetry via prolonged structural plasticity. **(a)** Interactions between two classes of AB pioneer neurons initiate lateralized axon remodeling. Different levels of Fas2 expression in SA^uni^ (low) versus SA^bi^ (high) support class-specific, NetB-mediated axon retraction (partial or complete for SA^bi^ or SA^uni^, respectively). **(b)** Following the completion of AB afferent lateralization, SA^uni^ and SA^bi^ switch their relative levels of Fas2 expression to high or low, respectively, accompanied by the bilateral dendrite ingrowth of FB relay neurons vΔA and SA^FB^. **(c)** The unilateral localization of Fas2-positive SA^uni^ axon terminals results in side-specific AB micro-circuits, with dominant afferent-afferent connectivity in AB-R versus afferent-efferent connectivity in AB-L.

Several findings indicate an active role of SA^uni^ in synaptic recruitment. First, lineage-related SA^bi^ and SA^FB^ form a large number of synapses with SA^uni^ in AB-R, whereas such afferent-afferent connections are almost absent in AB-L. Second, during development, vΔA first forms denser innervations in AB-L before invading the space occupied by Fas2-positive SA^uni^ in AB-R. This L-R difference is lost in brains with bilateral SA^uni^ innervation. Third, lineage-related LCNOpm neurons stochastically extend branches towards AB-R and in rare cases form a substantial number of synapses, as in the brain imaged for the Hemibrain dataset^18^. Both axonal pre-patterning of the AB primordia as well as secondary FB dendrite recruitment reveals a surprising amount of structural plasticity in conserved CX circuitry up to mid-pupa development. While fine-tuning of branching and synaptic growth also occur in second half of pupal development^30,34^, pattern of neuropile connectivity for bilaterally symmetric central brain circuits has already been finalized (with the exception of adult circadian plasticity^35^). As physical proximity in the transition from growth to connectivity is a strong predictor of synaptic partner choice^36^, a delay in the retraction from AB-L would allow SA^uni^ axons to recognize SA^bi^ processes as synaptic partners and thereby prevent their lateralized retraction.

Typically, neuronal remodeling of major neuronal circuits in *D. melanogaster* occurs synchronized in the central and peripheral nervous systems and is regulated by ecdysone signaling (reviewed in ^37^). The initial pruning is completed by 20 h APF^37^, i.e. before the AB primordium is formed. While ecdysone signaling was shown to be involved in establishing the left-right axis in the brain, SA^uni^ remodeling is initiated independently of ecdysone as lateralization is unaffected by the expression of a dominant-negative allele of the receptor EcR in AB neurons^17^.

In close proximity to the developing AB neuropil, FB neurons establish a commissural primordium in 1^st^ instar larva consisting undifferentiated neurites from embryonal cells^15^. The later ingrowth of secondary axons from later born neurons results in the maturation to the adult neuropil^15^. This sequence is reversed in AB development, where late-born tangential neurons form the primordium and establish the lateralized blueprint in early pupal development, before early-born plastic columnar neurons are recruited secondarily. During CX development, projections of columnar neurons establish arborizations in FB, PB, EB and NO by 28 h APF^38^. Pontine neurons form arborizations in the ventral FB later but are established by 40h APF^38^, a time point when vΔA starts invading the AB. Considering that roughly one third of vΔA neurons develop into pontine neurons and do not extend dendritic branches into the AB neuropils, the columnar vΔA projection pattern could have evolved from modified pontine neurons which gained responsiveness to AB derived signals. Delayed neuronal differentiation and synaptogenesis have been identified as key features of human cortex development supporting the generation of novel cell types and circuit motifs (reviewed in ^39,40^). In the *Drosophila* CX, the maintained responsiveness of intrinsically left-right symmetric neurons to guidance cues enables a small number of neurons to introduce a substantial level of asymmetry into the circuit. So far, the AB has only been described as a separate neuropil in *Diptera.* The physical segregation from the ventral FB might facilitate the asymmetrical pre-patterning without affecting the broader CX organization.

We found the *Drosophila* NCAM-homologue Fas2 to support both the asymmetric pre-patterning of the AB neuropils and the synaptic integration of the mature AB circuitry. Cell-adhesive molecules (CAMs) are critical factors during circuit assembly and remodeling, specifically their spatiotemporal control^41^. AB neuropils and afferent synapses do form in absence of Fas2, indicating that Fas2 function in axon guidance^28,42^ and synaptic elimination^28^ is not critical. In line with involvement of Fas2 in synapse formation, growth and stabilization^28,43,44^ we found support for a traditional role of Fas2 in SA^uni^ promoting synaptic growth indirectly measured by AB innervation area. However, we did not detect any cell intrinsic effect of Fas2 expression level in SA^uni^ on its neurite lateralization. A simple model in which Fas2-mediated trans-adhesion must be downregulated in AB-L to allow retraction of SA^uni^ axons, analogous to the remodeling of the mushroom body γ-lobe in which Fas2 must be downregulated to allow infiltration of astrocyte-like glia for pruning^29,45^, finds no support in our data. On the contrary, our data is most consistent with the assumption that during remodeling of SA^uni^, Fas2 functions non-cell-autonomously and non-adhesively to promote afferent lateralization. In SA^bi^ neurons, Fas2 appears to function as a signaling molecule. This is supported by the finding that both extracellular and intracellular domains are required to recapitulate the reduction in AB lateralization observed in the Fas2^PEST+^ overexpression experiments. Although the intracellular domain of Fas2 was shown to be involved in remodeling processes^29^ and synaptic growth^46^, Fas2’s intracellular domain is poorly described. Direct binding partners to its intracellular PDZ domaine have not been described to our knowledge. An indirect dependence on the intracellular domain has been shown for the regulation of Fas2 membrane availability by JNK signaling^29^ and the interaction with Appl to promote synaptic growth^46^. Furthermore, Fas2 expression can act as a non-cell autonomous repressor of EGFR during retinal differentiation^47^ and imaginal disc growth via heterophilic binding to Fipi and Elff, while promoting EGFR signaling by homophilic interactions^48^. Recently, a signaling function for Fas2 was also postulated independently of membrane anchoring^21^. Interaction with EGFR signaling and its role in synaptic stabilization^34^ seem especially consistent with our results indicating a role of Fas2 in modulating afferent-afferent synaptic connections.

A strong correlation between brain lateralization and cognitive performance has been recognized in humans and other animals. How the interplay between genetic programs and environmental factors influences adult brain lateralization is poorly understood. A reduction in bilateral asymmetry of SA^uni^ neurons has been shown to affect different aspects of *Drosophila* memory formation. Here we show that the genetic manipulation of a bilateral sister neuron of SA^uni^ results in cognitive performance in the context of appetitive olfactory memory and feeding-related decision making. This further increases the already substantial number of affected cognitive functions^6,8,32,49^. Interestingly, the degree of lateralization is impacted by environmental factors. The dynamic transient growth state before and during remodeling appears to be sensitive to developmental temperature. Low temperature led to lower lateralization in general by increasing the frequency of the fluctuating symmetrical SA^uni^ phenotype. We interpreted this as an effect of the increase of ectopic synaptic connections in slowed development^30^ and as evidence that neuronal proximity determines asymmetric synaptogenesis. However, it has been shown that the higher number of synaptic connections established at low developmental temperatures may be adaptive^30^.

In summary, our results demonstrate that CX lateralization initially arises through a robust genetic program, involving two classes of sequentially-born pioneer neurons which segregate their axons from a commissural system via interhemispheric cellular interactions. The non-adhesive and non-cell-autonomous function of Fas2 supports lateralized remodeling, a process that is notably sensitive to environmental factors. The secondary recruitment of processes from bilaterally symmetric CX neurons is intrinsically more variable and depends on the initial degree of afferent asymmetry as well as more dynamic cell-cell interactions thereby resulting in inter-individual variations.

## Methods

### *Drosophila* strains and genetics

Flies were kept on standard food medium. Following driver lines were used in this study: R72A10-Gal4^50^ (BDSC #48306), R72A10-lexA (BDSC #54191)^51^, R52H03-Gal4^50^ (BDSC #38849), R11F10-Gal4^50^ (BDSC no longer available), R66A02-Gal4^50^ (BDSC #39384), R66A02-lexA (BDSC #52688), R16E08-Gal4^50^ (BDSC #39416), R70H05-Gal4^50^ (BDSC #39554), R38D01-Gal4^50^ (#49996), R14B01-Gal4^50^ (BDSC #48597), R14B01-lexA (BDSC #52467)^51^, SS50477^10^ (BDSC no longer available), SS47384^10^ (BDSC #86590), SS47392^10^ (BDSC no longer available), Fas2-Gal4^Mz507^-Gal4 (B. Altenhein), Fas2^MiMIC12989^-Gal4^52^ (BDSC #77831). For neuronal labeling and clonal analysis we used MCFO^53^ (MCFO, BDSC #64086), Flybow 1.1B^54^ (BDSC #56802), 13x-lexAop-mCD8::GFP (Chr.II) (unkown source), 13x-lexAop-mCD8::GFP (Chr.III, BDSC #32203). 10x-UAS-mCD8::GFP (BDSC #32186), 10xUAS-MCD8::RFP & 13x-lexAop-mCD8::GFP (BDSC #32229), UAS-mCD8::cherry (BDSC #27392) and 10xUAS-FLAG (BDSC #62147). For experimental manipulation of Fas2 expression we used: UAS-Fas2^RNAi^ (BDSC #28990), UAS-Fas2^PEST+55^, UAS-Fas2^PEST-55^, UAS-intra-Fas2^PEST-^::YFP^27^ (A. Nose), UAS-extra-Fas2^PEST-^::YFP^27^ (A.Nose). Induced apoptosis was performed with UAS-hid^14^ (J.R. Nambu). Nrg^849^ (BDSC #35827) was used to prevent the development of interhemispheric comissures. We used the following constructs to analyse synaptic configuration and connectivity: UAS-syt::GFP (BDSC #6925), UAS-DenMark::cherry (BDSC #33062), UAS-brp::GFP (BDSC #36291), trans-Tango^56^ (BDSC # 77124), nysb-GRASP^57^ (BDSC #64315) and t-GRASP^58^ (BDSC #79039). Empty-Gal4 was created by insertion of the empty pBPGUw (addgene) vector following the protocol described by Pfeiffer et al.^59^.

### Connectome analysis

We accessed the hemibrain EM dataset^18^ (version 1.2.1) for connectome analysis via the neuPrint+^19^ browser interface (https://neuprint.janelia.org). neuPrint+ output from custom queries was downloaded and processed in R^60^ (v4.4.1, tidyverse^61^ v2.0.0).

### Nomenclature

In the hemibrain dataset, the SLP-AB^10^ category is divided into three neuronal categories: SA1, SA2, and SA3. We wanted to stick as closely as possible to the hemibrain nomenclature, but emphasized that SA1 and SA2 are the corresponding right and left hemispheric morphs of the same cell type, whereas SA3 is a separate type. Therefore, we have summarized SA1 and SA2 as SA^uni^ (after their typical unilateral AB-R projections) and renamed SA3 to SA^bi^ (for their bilateral projection pattern to AB-R and AB-L). SA^uni^ has also been referred to as Janus neurons^32^ and H neurons^8^ in publications. We have also named SAF SA^FB^ in this study to emphasize its similarity to the other afferents, but to refer to its specific projection to the dorsal FB.

### RNAi and overexpression experiments

UAS-construct flies were crossed with Gal4 driver lines and crosses were kept at 25°C or 18°C. As a control for Rubin lab Gal4 lines^50^, we crossed empty-Gal4 with the same UAS construct to maintain the genetic background in the controls, whereas in experiments with Fas2^Mz507^-Gal4, we crossed both Gal4 and UAS with Canton S. We evaluated the SA^uni^ projection pattern using immunodetection of R72A10-lexA driving lexAop-mCD8::GFP. The projection phenotypes were divided into two categories: unilateral (innervations only detectable in AB-R) and bilateral (innervations in AB-R and AB-L). If the row and column totals were greater than 5, a chi-squared test was performed. Otherwise, the Fisher exact test was used to test for statistically significant effects. We corrected for multiple comparisons using the Bonferroni method. All statistics were performed in R^60^ (v4.4.1). Plots were created with R ggplot2 v3.5.1^62^).

### Innervation volume and area quantification

To measure the 3D volume, surfaces were created in Imaris from fluorescence signals (GFP or cherry) by defining a region of interest, with identical settings for AB-R and AB-L surfaces. The acquired volumes were recorded and statistically analysed in Prism (GraphPad, v10.2.3). Statistical significance was tested using the Wilcoxon paired-samples test for consistent differences between AB-R and AB-L measurements within individuals. The Mann-Whitney U test was performed to compare grouped AB-R with grouped AB-L values.

Fiji^63^ (v2.14.0/1.54f) freehand tool was used to select the biggest dorso-ventral cross-section of the AB (R72A10-lexA>lexAop-GFP signal) and the EB neuropils (CadN signal). The sum of AB-R and AB-L and difference between AB-R and AB-L were normalized by dividing by EB cross section to account for variation in brain size between individuals. The normalized sum indicates general synaptic input to CX via AB innervations by afferent neurons ([AB-R+AB-L]/EB), normalized difference indicates direction and strength of lateralization of synaptic input right versus left ([AB-R-AB-L]/EB).

Effect sizes were compared computing rank-biserial correlation (R^60^ v4.4.1, effectsize^64^ v0.8.9), plots created in R (ggplot2^62^ v3.5.1 & beeswarm^65^ v0.4.0). Statistical analysis was done in R using Wilcoxon rank sum tests with continuity correction to compare Fas2^Mz507^-Gal4 data with controls, Kruskal-Wallis tests and Dunn post-hoc (rstatix^66^ v0.7.2) to compare more than 2 groups.

### Clonal analysis

MultiColor FlpOut or Flybow 1.1B was crossed with Gal4-driver lines and maintained at 25°C. Offspring were heatshocked in a water bath at 37°C (10 min for MCFO, 1 h for Flybow) at the late 3^rd^ instar stage.

### Analysis of pupal development

Pupal development was initially determined precisely to hours after pupal formation. These data were supplemented by pupae dissected in the early stages of development and their developmental stage approximated after imaging (*). Crosses and collected pupa were kept at 25°C.

To accurately time the pupae, pupae were collected within the first 0.5 hours after pupation (P0), and the subsequent time was counted as the hours ‘after pupation’ (APF). If the time from 0-0.5 hours before collection and the time for dissection to fixation are included, the h APF are accurate to within one hour. In order to approximate the h APF, pupae were dissected after head evagination until before eye pigmentation, and the progress of EB development was used to assess the stage of development. The EB primordia develop in both hemispheres before fusing into a single rod-shaped neuropile at the midline in early pupal development. The edges of this structure begin to curve ventrally at 12h APF and gradually form the adult ring structure. We calculated the percentage of ring closure: the angle between the ventral edge of the gap and the dorsal edge of the EB canal was subtracted from the 360° of the completed adult ring and divided by 360° (closure(%)=[360°-gap°]/360°). We used the ring closure to estimate the h APF based on the ring closure data of our precisely staged pupae. Approximation of h APF was based on linear regression (Fig. S8, Prism v10.2.3). EB closure and the portion of brains with segregated AB precursors in precisely staged pupae was plotted in Prism over h APF. For more information see supplementary material.

### Confocal acquisition and image analysis

Confocal imaging was performed using a Leica TCS SP5II microscope with a 20X oil immersion and a 40x water immersion objective. Images were processed with ImageJ and Imaris (Bitplane, v9.3.1).

Images, especially for SA^uni^ phenotype counting, were acquired at 400 Hz with a resolution of 512x512. Representative morphological data were acquired at 200 Hz and a resolution of 1024x1024.

### Immunohistochemistry

Brains were dissected in PBS buffer and fixed in 4% PFA in PBS for 25 min at room temperature. Samples were washed 4 times in 0.3% Triton-X in PBS before blocking for 1h in 10% goat serum in PBS. Incubation with primary antibodies was performed at 4°C overnight. Samples were washed 4 times in 0.3% Triton-X in PBS followed by incubation with secondary antibodies overnight at 4°C. Samples were washed 4 times in 0.3% Triton-X in PBS before mounting in VECTASHIELD (Vector Laboratories) medium.

Primary antibodies used for this study were rat anti–CadN extracellular domain [cells from DN-Ex no.8, 1:10; Developmental Studies Hybridoma Bank (DSHB), in-house production], mouse anti-Fas2 (1D4) intracellular domain [1:10, cells from DSHB, in-house production], rabbit anti-GFP (1:1000; Invitrogen), rabbit anti-HA and rat anti-FLAG. Secondary antibodies used for this study were goat anti-rabbit Alexa 488 (1:500, Invitrogen), goat anti-mouse highly-cross absorbed Alexa 488 (1:500, Invitrogen), goat anti-mouse highly-cross absorbed Alexa 568 (1:300, Invitrogen), goat anti-rabbit Alexa 568 (1:300, Invitrogen) and goat anti-rat Alexa 647 (1:500, Invitrogen). All antibodies were diluted in 10 % goat serum in PBS at the indicated ratios.

### Satiety-dependent fructose drive

The flies were fasted for 30 hours at 29°C on 1% agarose gel. All flies were tested individually in the FlyPad^67^ apparatus (Easy Behavior) following the protocol described by Musso et al.^32^.The two electrodes were prepared with 3 µL 1% agarose in 50 mmol/L sugar solutions, one electrode with glucose, the other with fructose. During the experiments, the interactions of the flies with the presented sugar were recorded at each electrode for 1 hour. Preference indices were calculated from the interactions (PI = [fructose-glucose] / [all interactions]). The FlyPad output was preprocessed for further analysis in R. The results were plotted and statistically analyzed in Prism. The Wilcoxon paired-samples test was performed to test for consistent differences in the number of glucose/fructose interactions between individuals, and the Mann-Whitney U test was used to test for statistically significant effects between the performance indices per group.

### Single-Session Appetitive Olfactory Conditioning

The flies tested were raised on standard cornmeal food and a 12–12 h light–dark cycle at 18°C. Fasting, training and testing was performed at 23°C and 60% humidity. The training and tests were carried out in an elevator T-maze apparatus^68^ following the protocol established by Krashes and Waddell^33^. The flies were fasted for 20 hours humidity on 1% agar before training. For the training, the flies were first exposed to the conditioned stimulus - (CS-) without sugar reward (filter paper soaked in water) for 1 min. Then the conditioned stimulus + (CS+) paired with a sugar reward (saturated sucrose solution on filter paper) was presented to the flies for 1 min. The odorants 3-octanol (OCT) or 4-methylcyclohexanol (MCH) were used alternately as CS+ or CS- in successive test runs with the same genotype. To prepare odor solutions, 8 µL OCT or 13 µL MCH were diluted in 8 mL mineral oil. To test immediate memory performance, flies were directly transferred into the T-maze after training. To test 24h memory expression, flies were kept on 1% agar for 24h after training. Testing was performed in complete darkness. The flies had 2 minutes to choose either the arm with the CS+ or the arm with the CS-. For each run, an index was calculated by dividing the difference between the number of flies in the CS+ arm and the number of flies in the CS arm of the T-maze by the total number of flies in both arms. The performance indices (PIs) were calculated as the mean value of the indices for the reciprocal CS+/CS- training runs. Results were plotted and tested for statistically significant effects using Prism.

## Supporting information

Supplemental Information 1 - Statistics exact values

## Acknowledgments

We would like to thank Iris Salecker (Flybow), Benjamin Altenhein (Fas2-Gal4^Mz507^), Akahiro Nose (UAS- intra- & extra-Fas2::YFP), and Corey S. Goodman (UAS-Fas2PEST+/-), as well as the Bloomington Stock Center for providing materials and fly stocks. We thank Scott Waddell and his lab, especially Bhagyashree Senapati, for providing the opportunity to conduct memory experiments at the CNCB, University of Oxford, and for their supervision and discussions during this period. We also wish to thank Wolfgang Kallina, Sigrid Ilgerl, Daniela Bartel, Alexandra Grimm, and Alena Litin for their technical support, and the Hummel Lab for stimulating discussions and their critical comments on the manuscript. We acknowledge the early exploratory work of Amadeo Mattia, Stefan Trkulja, Charlotte Schönherr, Sonja Bogner, Barbara Simpson, Laurin Tomasek, Heidi Roth, Hana Vokač, Rosa Gredler, and Felix Kapelari, Teodora Kolarova, Cornelia Ignitsch, Áron Bautista-Soldevila and Mariam Kassem. Funding: University of Vienna, CoBeNe Doctoral School (uni:docs fellowship), Cluster of Excellence "Neuronal Circuits in Health and Disease" (Fonds zur Förderung der wissenschaftlichen Forschung, FWF).

## Author Contributions

J.M. conceptualized and investigated the study, acquired and analyzed data, interpreted the results, and wrote the article. D.M., A.JG., R.K., Z.A., S.K. and T.K. acquired and analyzed data. T.H. conceptualized the study, supervised the research, analyzed and interpreted the data, and acquired funding.

## Additional information

All data needed to evaluate the conclusions of this paper are present in the paper and/or the Supplementary Materials. Additional data related to this paper are available upon request.

---

**Fig S1.**
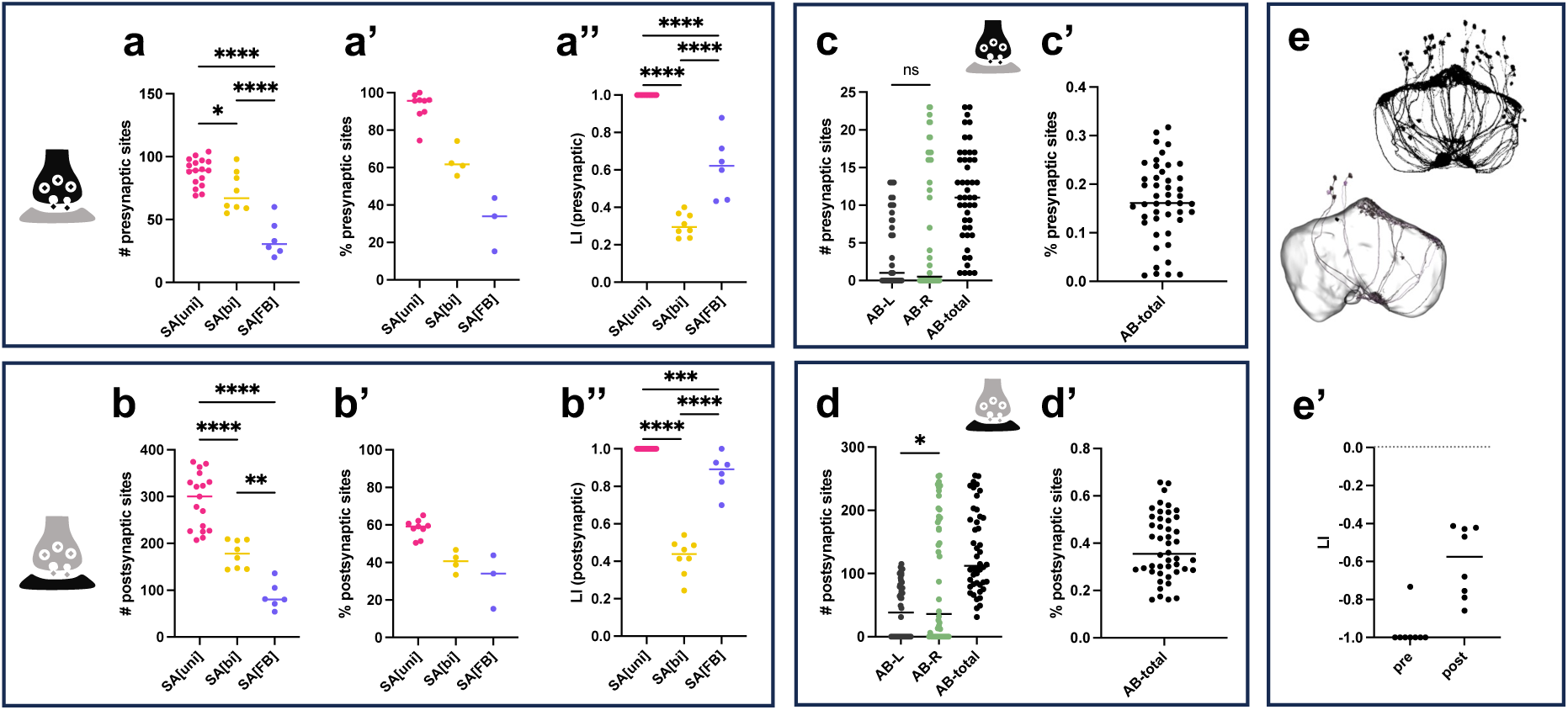
Bilateral asymmetry in in synaptic connectivity for AB neurons classes. **(a)** Number of presynaptic sites in AB-R and AB-L per neuron for the three main SA afferent neuron types. SA^uni^ neurons have the highest number of presynaptic sites, SA^FB^ the lowest, one-way ANOVA revealed a significant difference between neuronal types in number of presynaptic sites, F(2,28)=36.54, p=1.6*10^-8^. **(a’)** Percent of presynaptic sites in AB-L and AB-R compared to the total number of presynaptic sites of a neuron. While almost all presynaptic sites of SA^uni^ are located in the AB, SA^bi^ and SA^FB^ have a substantial amount of presynaptic sites in the SLP or the dorsal FB respectively. **(a’’)** Lateralization Index per neuron by afferent type. Positive Indices indicate % of total AB presynaptic site that are more on the right (AB-R – AB-L)/(AB-all). All afferents show a right-directed lateralization of presynaptic site. SA^bi^ is the least lateralized afferent. one-way ANOVA revealed a significant difference between neuronal types in presynaptic lateralization (LI), F(2,28)=213.7,p=1.1*10^-17^. **(b)** Number of postsynaptic sites per neuron in AB-R and AB-L for the three main SA afferent neuron types. One-way ANOVA indicated a significant difference between neuronal type in number of postsynaptic sites, F(2,28)=43.01, p=2.9*10^-9^. **(b’)** Percent of postsynaptic sites in AB-L and AB-R compared to the total number of presynaptic sites of a neuron. All “afferents” receive substantial input in the AB neuropiles, SA^uni^ receives more input in AB-R than from the SLP. **(b’’)** Lateralization Index per neuron by afferent type. Positive Indices indicate % of total AB postsynaptic site that are more on the right (AB-R – AB-L)/(AB-all). Suprisingly, AB input to SA^FB^ comes almost exclusively from AB-R (One-way Anova, F(2,28)=218.2,p=8.4*10^-18^). **( c, d)** Numbers of synaptic connections by vΔA in AB-L and AB-R and **(c’, d’)** the % of total synaptic connections located in the ABs per vΔA neuron. Mann-Whitney U test reveals significant difference in number of postsynaptic sites of vΔA neurons in AB-L compared to AB-R, U= 780, p=0.0241, but indicates that that numbers of presynaptic sites of vΔA neurons are most compatible with the hypothesis they are not differently distributed between AB-L and AB-R, U= 916, p=0.2397. **(e)** Some vΔA neurons bilaterally innervate AB-L and AB-R **(e’)** Normalized difference in pre- and postsynaptic sites between AB-R and AB- L in bilaterally projecting vΔA neurons (Lateralization Index, (R-L)/(R+L)) indicates mostly postsynaptic sites in AB-L. SA^uni^ in magenta, SA^bi^ in yellow and SA^FB^ in blue. Bars indicate group medians. Significance threshold are: *<0.05, **<0.01, ***<0.001, ****<0.0001. Pairwise comparisons by Tukey HSD test. For B’ and C’ only neurons of the right hemisphere were taken into account, because left SLP is not included in the hemibrain dataset.

**Fig S2.**
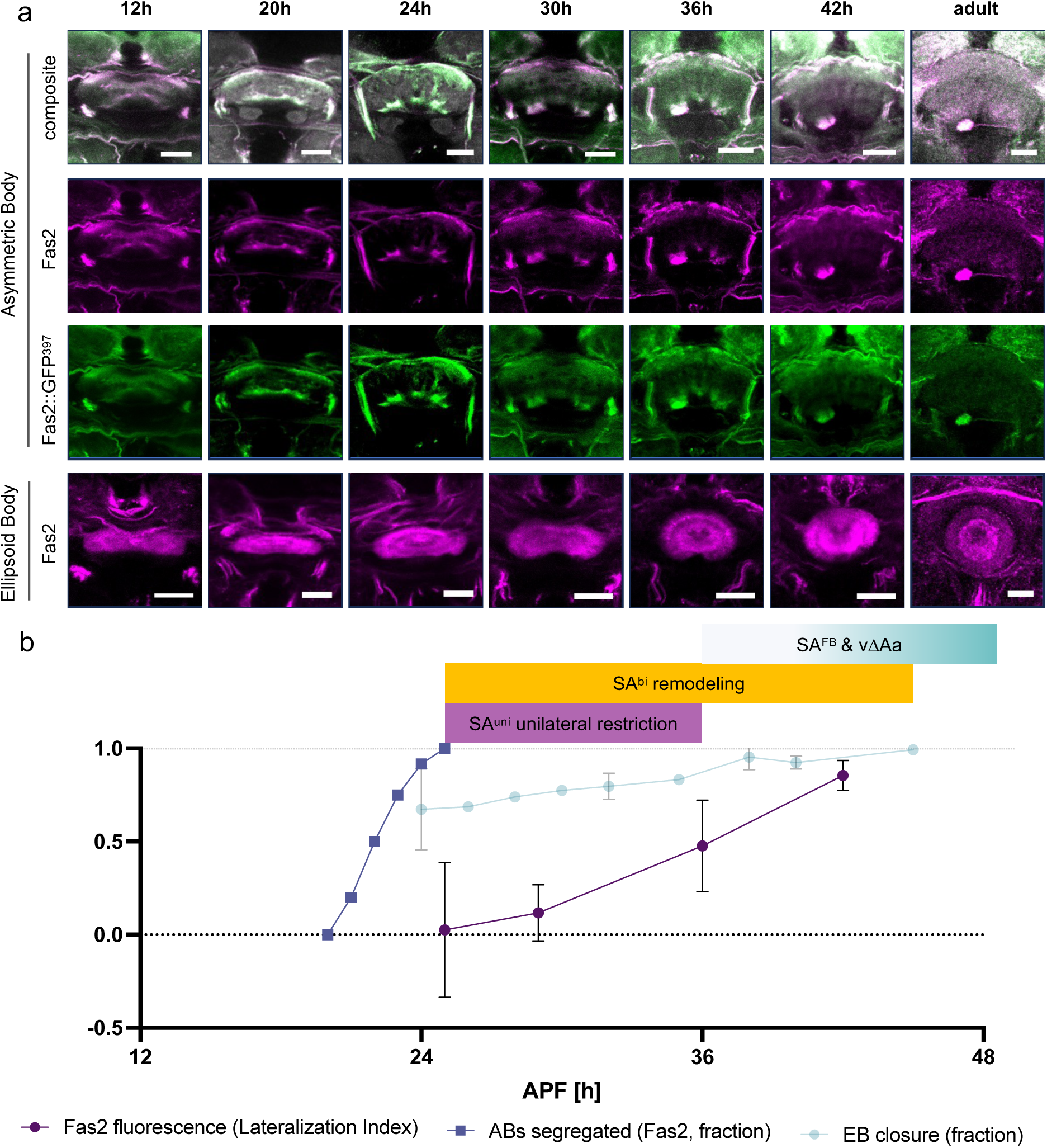
Fas2 is dynamically expressed in in AB development. **(a)** Pupa brains dissected at the indicated number of hours after puparium formation (h APF). Immunodetection of Fas2::GFP^397^ (green) and Fas2 (magenta) and CadN (grey) antibody staining. **(b)** Timeline of AB development based on Fas2 expression. From 21h to 25h APF Fas2-positive AB precursors segregate from an accumulation of Fas2 positive layers in the developing ventral FB (rate of analysed brains with segregated AB precursors, blue squares). At 25h APF signal from Fas2 immunodetection in AB-R and AB-L did not show a population-level directed lateralization, but detected fluorescence in AB-R became increasingly more intense compared to AB-L up to 42h APF (Lateralization Index (AB-R-AB-L)/(AB-R+AB-L), magenta dots (mean LI), bars indicate 95% Confidence Intervals). The maturation of the EB from an elongated oval to a circular shape occurs in the same time window of CX development, shows low variability and can be used as an approximation for the number of hours after pupal formation (fraction of closure measured by ventral gap(°)/360, light blue dots (mean fraction, bars indicate 95% confidence intervals). All scale bars, 20µm.

**Fig S3.**
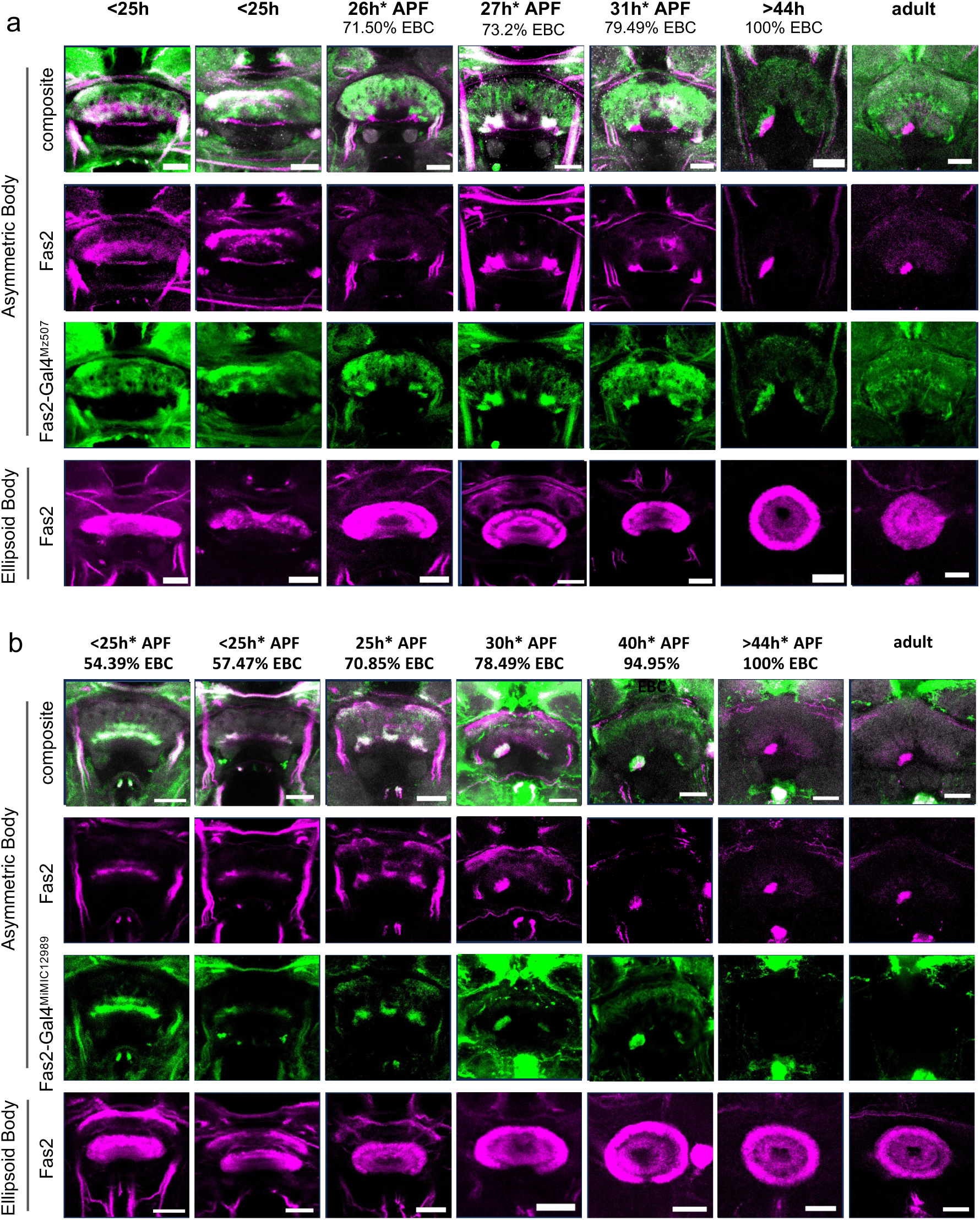
Expression of Gal4 under control of endogenous regulatory regions does not fully mimic Fas2 expression in the developing and adult AB. **(a)** Fas2-Gal4^MZ507^ expression is restricted to the dorsal AB-R and AB-L precursors. Bilateral expression is detectable until mid-pupal stage, when expression in the AB neuropils ceases altogether. Immunodetection of UAS-GFP driven by Fas2-Gal4^MZ507^ (green) and Fas2 (magenta) and CadN (grey) antibody staining. **(b)** Fas2-Gal4^MiMIC12989^ expression is restricted to the dorsal AB-precursors. Endogenous expression of Fas2 appeared unaffected. Gal4 expression is absent in the adult AB. Immunodetection of UAS-GFP driven by Fas2-Gal4^MiMIC12989^ (green) and Fas2 (magenta) and CadN (grey) antibody staining. All scale bars, 20µm.

**Fig S4.**
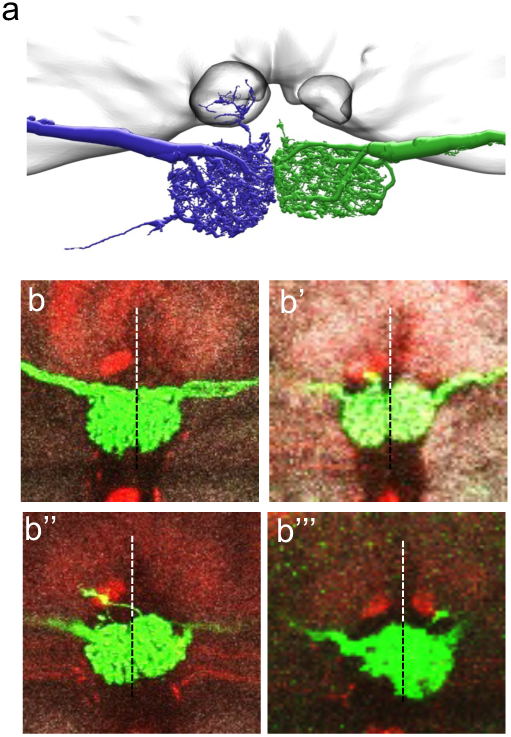
Sporadic AB innervation by LCNOpm neurons. **(a)** LCNOpm neurons innervate the ipsilateral Noduli in relation to their cell body position. The hemibrain shows a substantial number of postsynaptic connections from postsynaptic sites formed by right neurons in AB-R, but no synaptic connections for left hemisphere neurons in AB-R or AB-L. **(b-b’’’)** We found LCNOpm neurons to typically not innervate the AB neuropiles. In 17% of analysed brains right **(b’’)** or left **(b’’’)** LCNOpm neurons innervate in the ABs with a clear preference for targeting AB-R.

**Fig S5.**
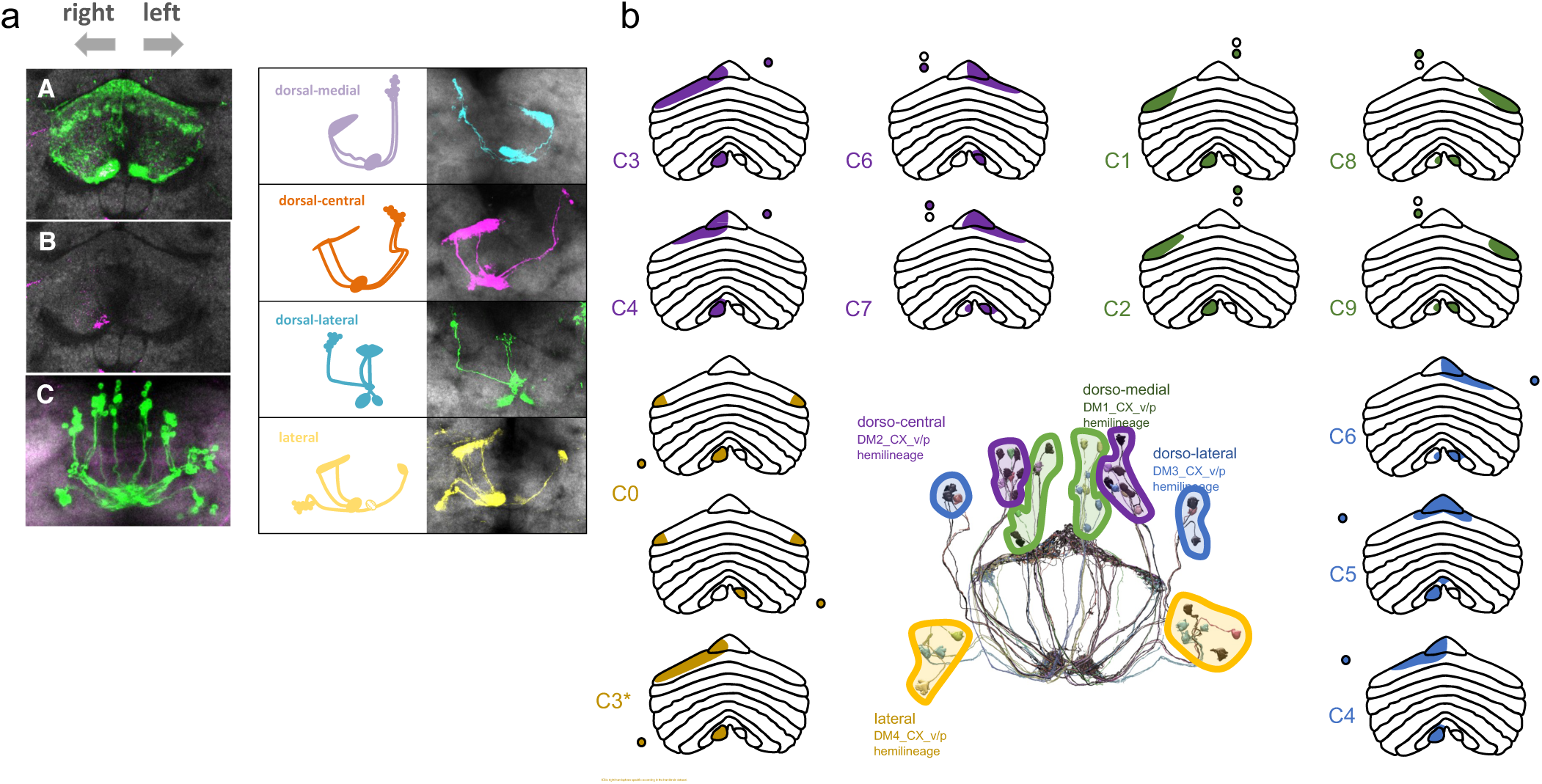
Adult vΔA neuronal morphology. **(a)** vDelta cell bodies are positioned in four clusters per hemisphere. Immunofluorescence detection from UAS-GFP (green, left panel) / UAS-Flybow1.1B (right, panel: cyan, magenta, green, yellow) driven by R70H05-Gal4 and Fas2 (left panel, magenta) and CadN antibody staining. **(b)** These cluster indicate their descendance from lineages DM1-4. Neurons of each cluster follow a stereotyped projection pattern.

**Fig S6.**
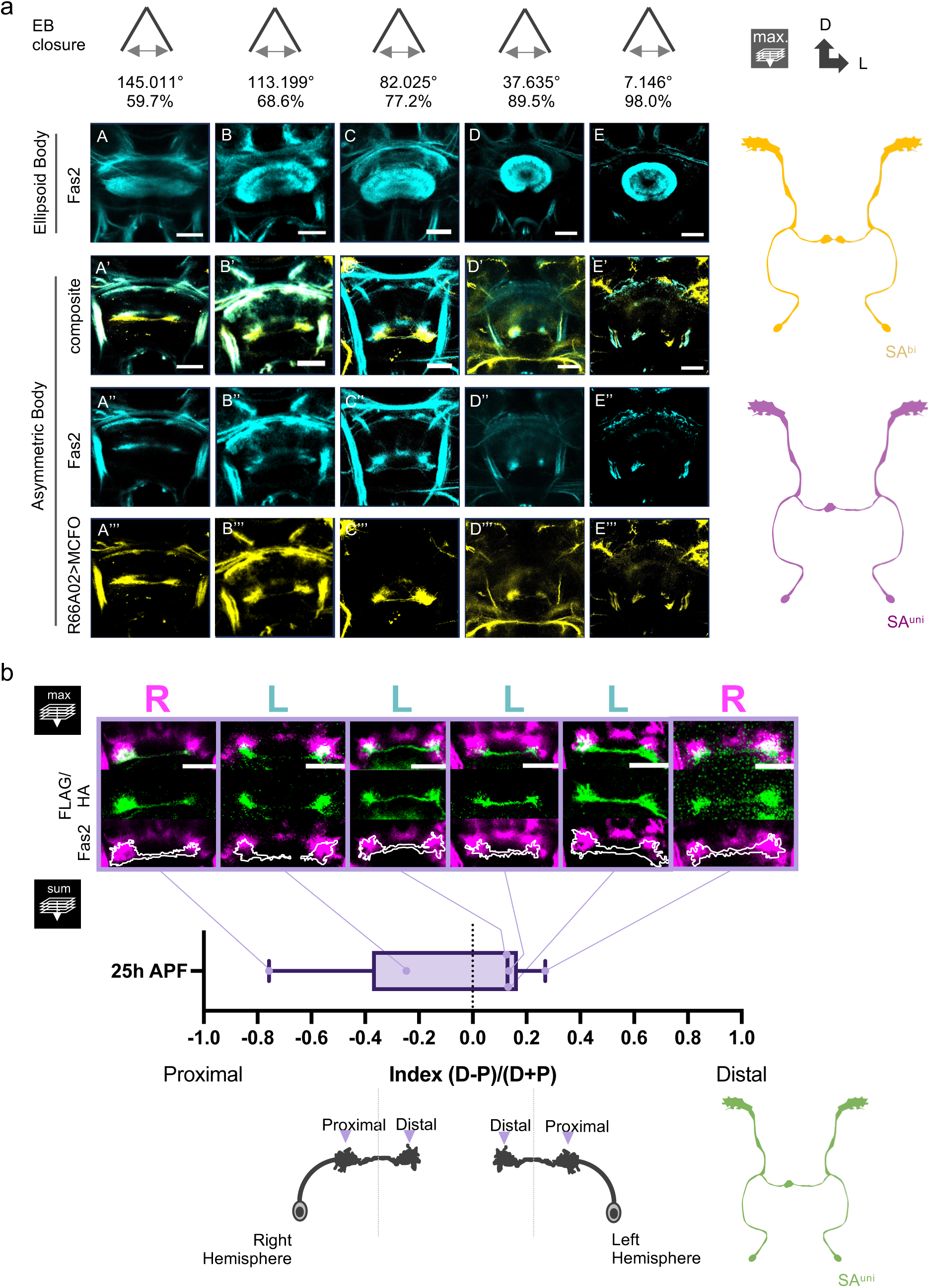
The SA^bi^ neurones densely innervate the dorsal AB primordium, SA^uni^ neurons the ventral primordium. **(a)** Clonal analysis of SA^bi^ during pupal developement based on immunodetection of UAS-Flag/HA (MCFO) driven by R66A02-Gal4 in hsFLP induced mosaics (yellow), and Fas2 antibody staining (cyan). EB gap closure (%) indicates progress of pupal development. **(b)** At 25h SA^uni^ neurons innervate the ventral AB-R and AB-L primordia. Measurements from immunodetection of GFP signal from single cell clones (Boxplot). No consistent difference in innervation strength between proximal and distal innervations of the same neuron were detectable, while we found a tendency for the distal innervations to be denser. Immunodetection of UAS-Flag/HA (MCFO) by R11F10-Gal4 in hsFLP induced mosaics (SA^bi^ clones in yellow, SA^uni^ clones in magenta) and Fas2 antibody staining (cyan). All scale bars, 20µm.

**Fig S7.**
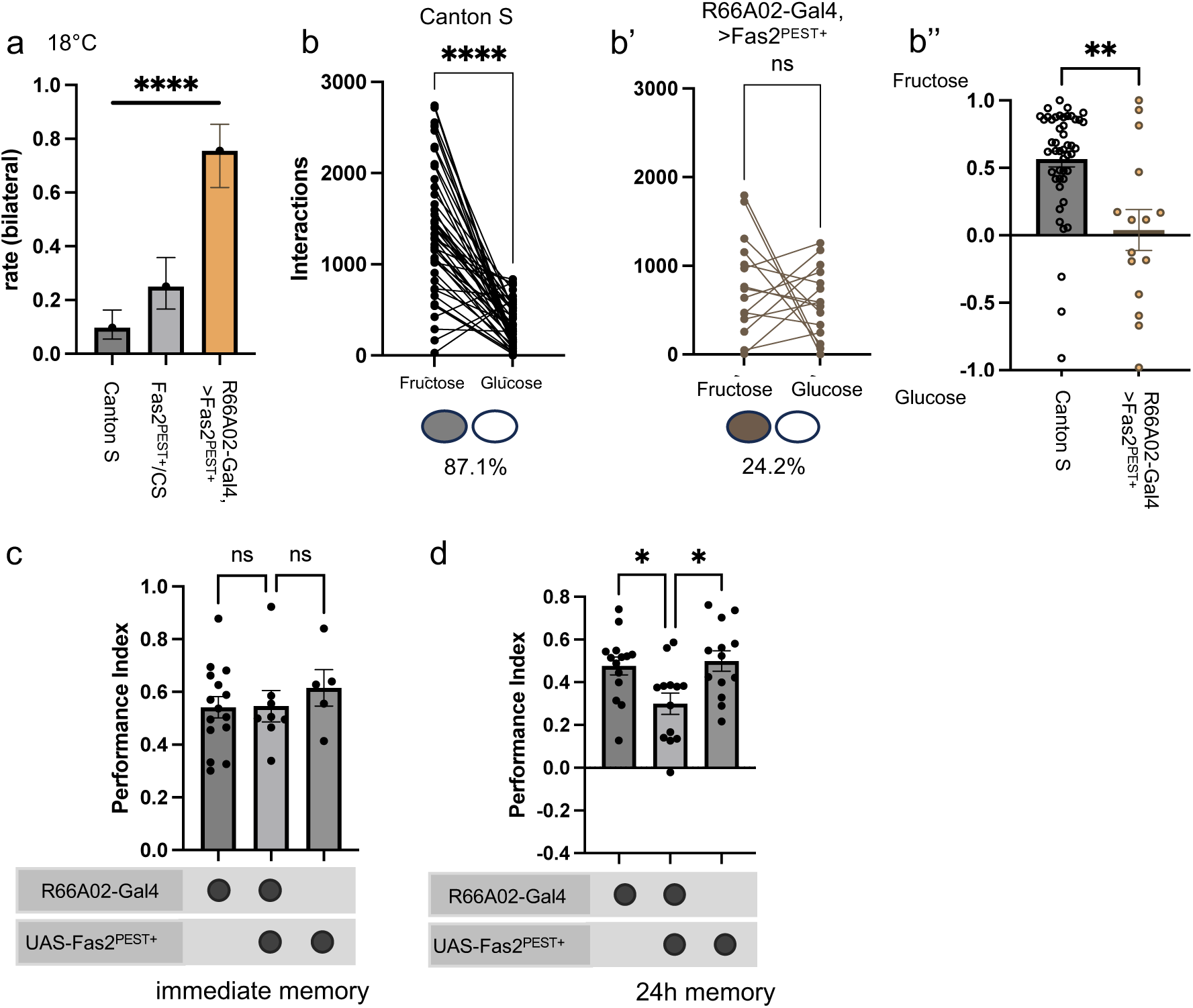
Bilateral SAuni projections induced by Fas2 change behavioural expression. **(a)** Penetrance of the SA^uni^ bilateral projection phenotype is statistically significantly different between flies expressing Fas2^PEST+^ driven by R66A02-Gal4 and control flies (Canton-S and UAS-Fas2^PEST+^, χ^2^). **(b,b’,b’’)** While wildtype flies (Canton-S) showed the described post-fasting preference for fructose over glucose, no consistent preference was detectable in flies expressing Fas2^PEST+^ driven by R66A02-Gal4 (Wilcoxon Test for paired samples; Mann-Whitney U). **(c)** Flies expressing Fas2^PEST+^ driven by R66A02-Gal4 express immediate memory of odour with a sugar reward, but **(d)** expression of sugar-reward memory after 24h is statistically significantly reduced compared to controls (Kruskal-Wallis/Dunn). Significance threshold are: *<0.05; **<0.01; ***<0.001; ****<0.001.

**Fig S8.**
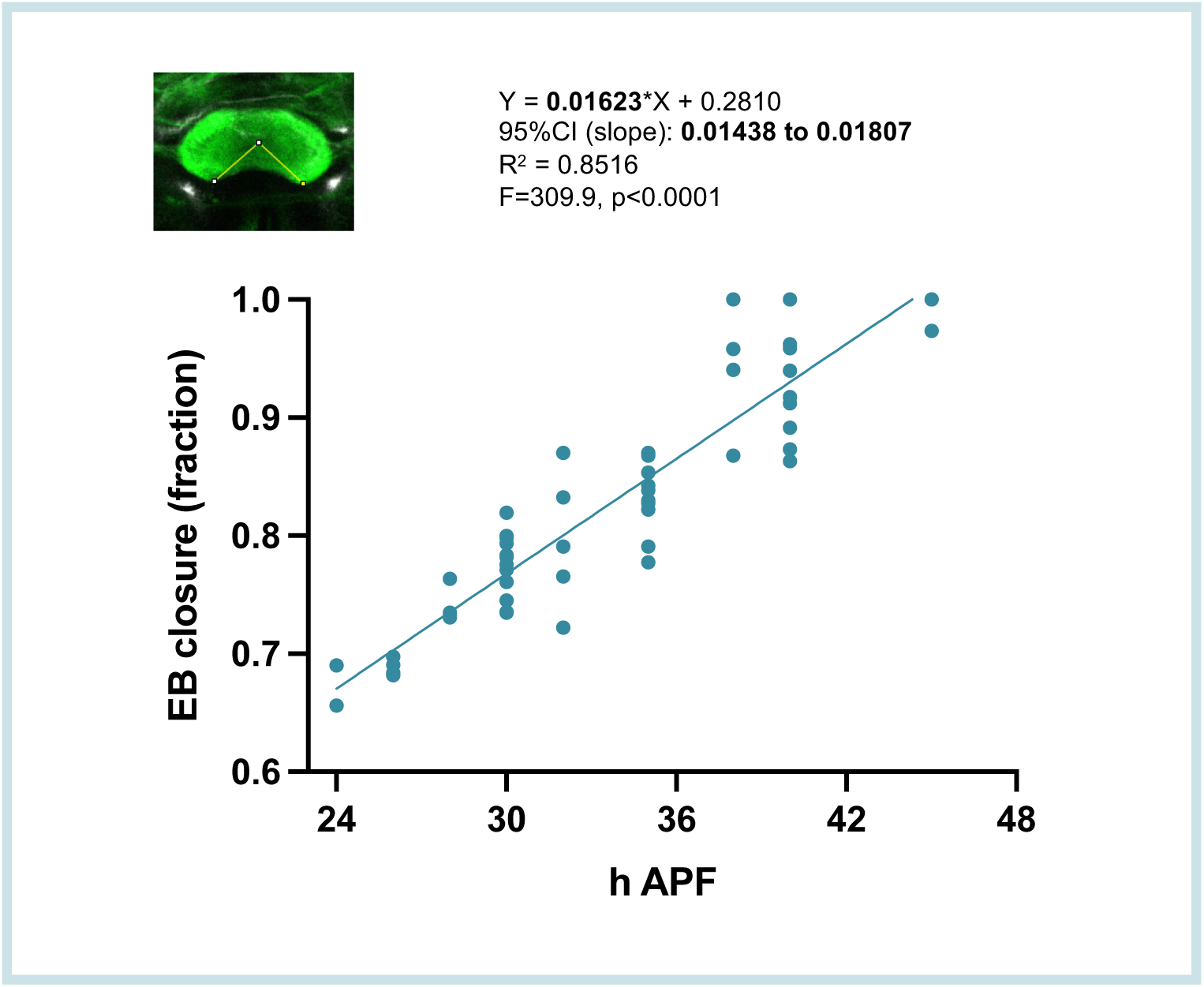
Correlation of EB closure and hours of pupal development.

